# Energetics of a protein disorder-order transition in small molecule recognition

**DOI:** 10.1101/2021.08.05.454052

**Authors:** Cesar Mendoza-Martinez, Michail Papadourakis, Salomé Llabrés, Arun A. Gupta, Paul N. Barlow, Julien Michel

**Affiliations:** EaStCHEM School of Chemistry, University of Edinburgh, David Brewster Road, Edinburgh, EH9 3FJ, United Kingdom

## Abstract

Many proteins recognise other proteins via mechanisms that involve the folding of intrinsically disordered regions upon complex formation. Here we investigate how the selectivity of a drug-like small molecule arises from its modulation of a protein disorder-to-order transition. Binding of the compound AM-7209 has been reported to confer order upon an intrinsically disordered ‘lid’ region of the oncoprotein MDM2. Calorimetric measurements revealed that truncation of the lid region of MDM2 increases the apparent dissociation constant of AM-7209 250-fold. By contrast, lid truncation has little effect on the binding of the ligand Nutlin-3a. Insights into these differential binding energetics were obtained via a complete thermodynamic analysis that featured adaptive absolute alchemical free energy of binding calculations with enhanced-sampling molecular dynamics simulations. The simulations reveal that in apo MDM2 the ordered lid state is energetically disfavoured. AM-7209, but not Nutlin-3a, shows a significant energetic preference for ordered lid conformations, thus shifting the balance towards ordering of the lid in the AM-7209/MDM2 complex. The methodology reported herein should facilitate broader targeting of intrinsically disordered regions in medicinal chemistry.

## Introduction

Over 25% of proteomes in eukaryotes consist of proteins that lack a well-defined tertiary structure.^1,2^ Nearly half of all proteins contain significant intrinsically disordered regions (IDRs), i.e. contiguous stretches of 20-50 amino acids that remain disordered in native conditions. IDRs are often key players in molecular recognition processes that underpin the affinity and selectivity of protein-protein interactions (PPIs). ^3^ A common scenario is that an IDR motif in a protein undergoes a disorder-to-order transition upon binding to another protein, leading to formation of a low-affinity high-selectivity complex.^4^ Classical examples of disorder-to-order transitions include PPIs where one or both partner fold(s) into a well-structured protein upon binding.^5–7^ However a growing number examples illustrate that one or both partner in a PPI may remain significantly disordered in the complex.^8–12^

Ample evidence points to the important roles of IDRs in the pathophysiology of diverse diseases such as cancers, diabetes or neurodegenerative disorders.^13^ Consequently there is strong interest in developing therapeutic agents that interact with IDRs to modulate protein-protein interactions.^14,15^ An appealing strategy would be to mimic Nature and identify small molecules that induce disorder-to-order transitions on binding to a target of interest.^16–18^ To date examples of such small molecules have been largely discovered by serendipity. Consequently there is an unmet need for methodologies that facilitate the design of modulators of disorder-to-order transition in protein structures.^19–26^

With the view of advancing general understanding of small molecule-IDR interactions, the present report focuses on elucidating the energetics of a disorder-to-order transition mechanism observed upon binding of the small molecule AM-7209 to the N-terminal domain of the MDM2 protein (Figure 1A,1B). MDM2 is a negative regulator of the tumor suppressor p53.^27,28^ MDM2 is a validated drug target that has attracted vigorous medicinal chemistry efforts.^29–31^ Several p53/MDM2 antagonists have entered clinical trials for diverse oncology indications.^32,33^ AM-7209 is a lead molecule from the piperidinone family of MDM2 ligands, that were serendipitously found to induce ordering of the N-terminal ‘lid’ IDR of MDM2 upon binding.^34,35^ By contrast the p53/MDM2 antagonist Nutlin-3a binds to MDM2 without ordering the lid region.^35,36^ Here we provide a rationale for the different binding mechanisms of AM-7209 and Nutlin-3a by combining calorimetric measurements with a novel method for performing adaptive absolute free energy of binding calculations, and with enhanced-sampling molecular dynamics (MD) simulations. The results pave the way for broader targeting of IDRs in structure-based drug design campaigns.

**Figure 1.**
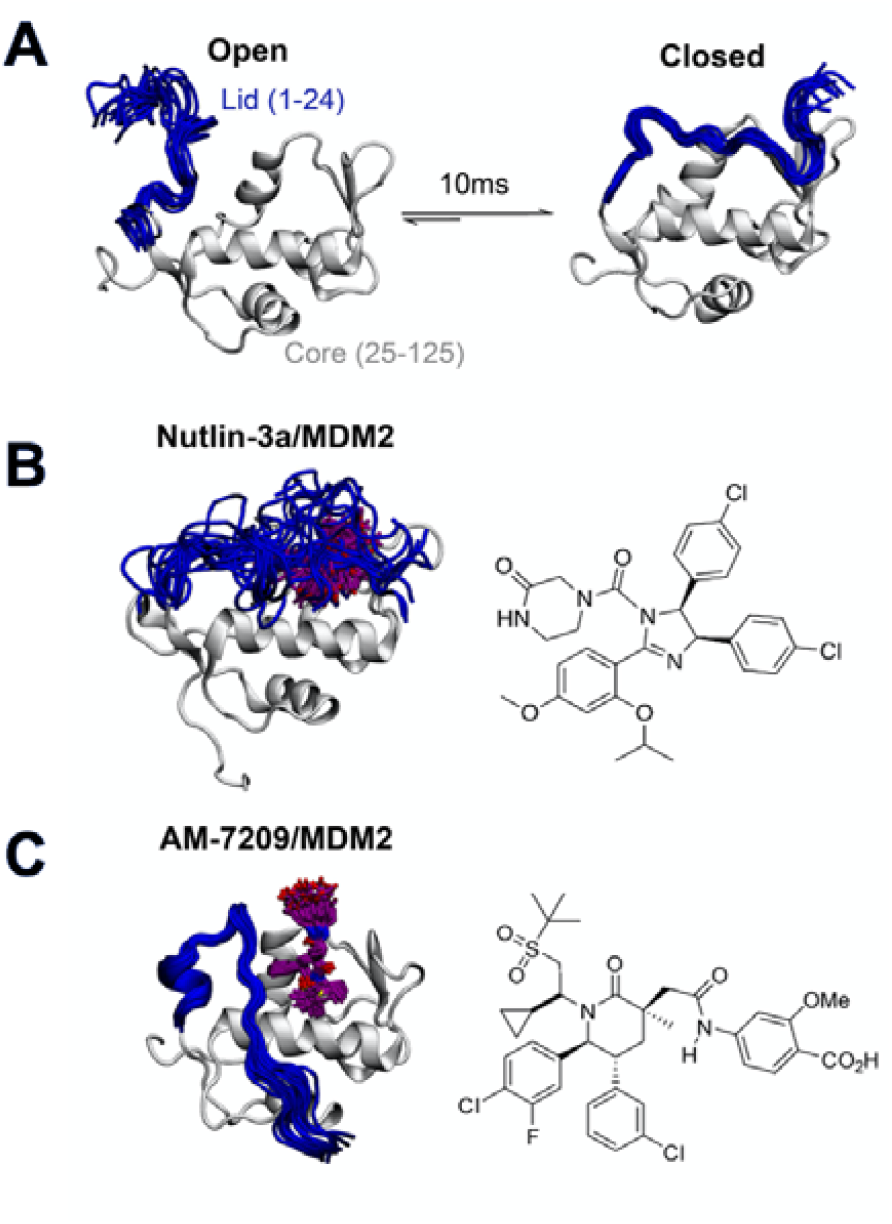
Ligand-specific modulation of conformational preferences of the MDM2 N-terminal domain lid IDR. **A)** In apo MDM2 the lid region is disordered and exchanges on a millisecond timescale between ‘open’ and ‘closed’ disordered conformational states that regulate access to the p53-binding site.^37,38^ **B)** Previous work from Bueren-Calabuig and Michel suggests the ligand Nutlin-3a binds MDM2 with the lid in ‘closed’ state broadly similar to that seen in apo.^39^ **C)** Binding of the MDM2 ligand AM-7209 orders the MDM2 lid region into a helix-turn-strand motif.^35,39^ Representative snapshots of the lid conformations are taken from the simulations reported in this manuscript.

## Results and discussion

### Truncation of the MDM2 lid IDR modulates AM-7209 affinity by 250-fold

We prepared protein constructs for human MDM2 6-125 and MDM2 17-125 (Figure S1, Figure S2, referred to as MDM2-Lid and MDM2-Lid/short from here on) and carried out ITC measurements to study the effect of lid truncation on Nutlin-3a and AM-7209 binding Previous reports indicate that both compounds bind to MDM2 with a 1:1 stoichiometry.^34,36^ The present experiments gave well behaved titrations with similar errors for all complexes (Figure S3, Figure S4, Table S1). In the case of Nutlin-3a lid truncation only weakly increases the apparent dissociation constant, by 1.2-2.3 fold (Fig 2A), without significant changes in enthalpy or entropy of binding (Fig 2B-C). These observations are in broad agreement with previous reports.^35^ Our measurements reveal that AM-7209 is an extraordinarily potent ligand for MDM2-Lid, with an apparent *K*_d_ value of ca. 5±1 pM (Fig 2A). In the case of AM-7209, lid truncation causes a significant increase in the enthalpy of binding, and a decrease in the entropy of binding (Fig 2B-C). Thus, changes in enthalpy favour binding of AM-7209 to MDM2-Lid over MDM2-Lid/short. This suggests that additional interactions between the MDM2 lid region and AM-7209 or the MDM2 core region are present in the AM-7209/MDM2-Lid complex in comparison with the AM-7209/MDM2-Lid/short complex. The lack of difference in enthalpies of binding of Nutlin-3a between the two MDM2 constructs suggest that AM-7209 induces a distinct conformational change in the MDM2 lid region over Nutlin-3a. As changes in entropy disfavour binding of AM-7209 to MDM2-Lid over MDM2-Lid/short, this suggests that such lid conformational change is associated with increased rigidity of the lid region in the AM-7209/MDM2-Lid complex over the Nutlin-3a/MDM2-Lid complex.

**Figure 2.**
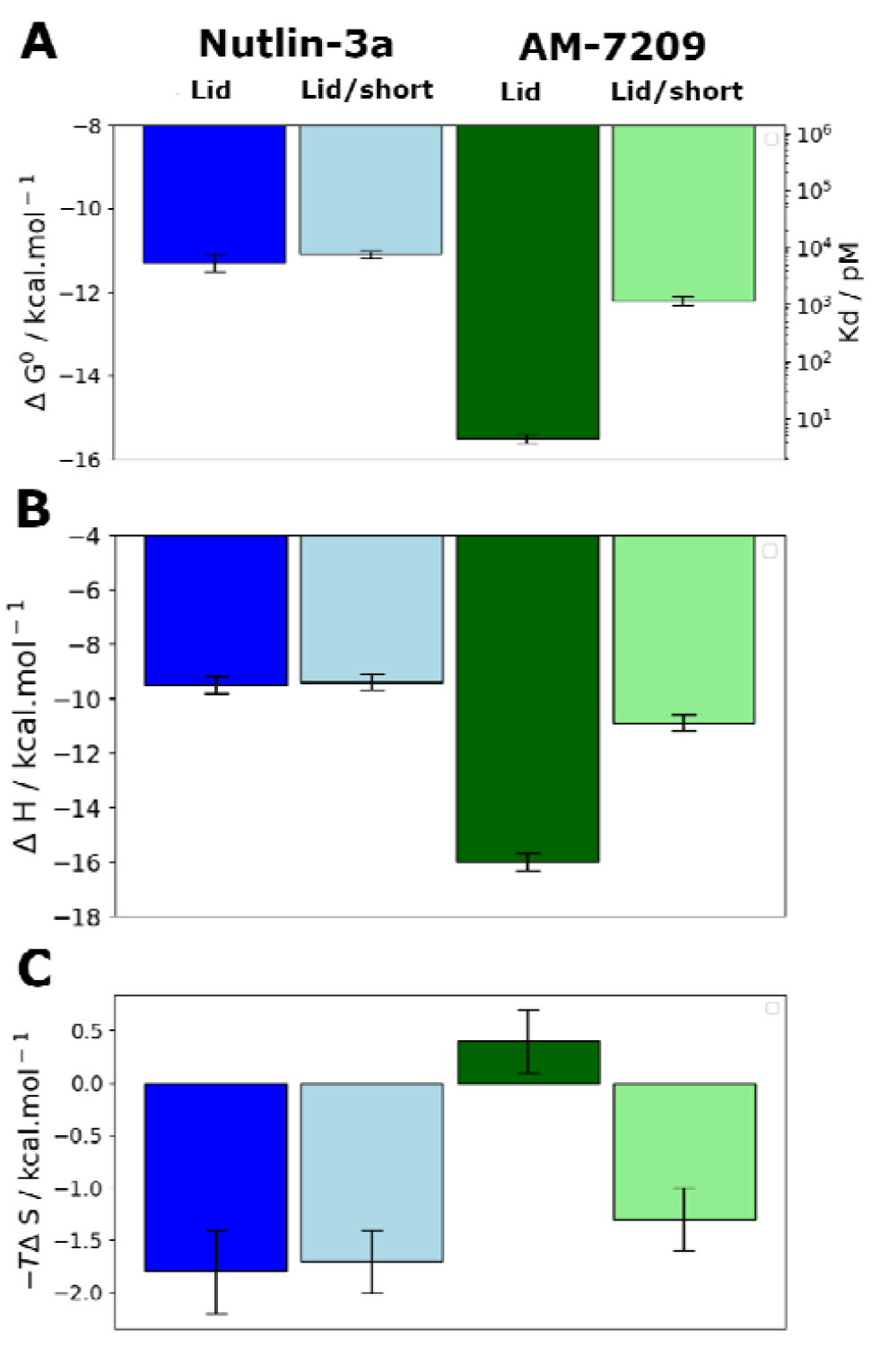
MDM2 lid truncation has a differential effect on thermodynamic signatures of Nutlin-3 and AM-7209 binding. **A)** Apparent standard Gibbs free energies of binding and dissociation constants measured by ITC experiments. **B)** Enthalpies of binding. **C)** Entropies of binding. Dark blue: Nutlin-3a/MDM2-Lid. Light blue: Nutlin-3a/MDM2-Lid/short. Dark green: AM-7209/MDM2-Lid. Light green: AM-7209/MDM2-Lid/short. Error bars denote ±1σ (*n*=3).

The net effect of lid truncation in the case of AM-7209 binding to MDM2 is a remarkable 250-fold increase in apparent *K*_d_.

### Spontaneous ordering of the MDM2 lid is energetically disfavoured in apo MDM2

Clarification of the intrinsic energetic preferences of the MDM2 lid IDR was sought by using molecular dynamics (MD) simulation methodologies. Forcefield development for molecular dynamics simulations of IDPs is an active research area.^25,40^ The choice of solvent model and protein forcefield greatly influences the conformational ensembles gathered from MD simulations.^41–43^ For the specific case of protein IDRs simulations, forcefields that correctly describe both the intrinsically disordered regions and the folded regions of the protein are required.^25,44^ There is no consensus yet on which is the best forcefield for IDRs as the accuracy of the predictions seem to depend on the studied protein. Here we used the amber99SB-ildn-nmr forcefield because it was shown to reproduce reasonably the behaviour of ordered motifs in proteins,^45^ and the disordered N-terminal tail region of histone H3.^46^ Previous studies with this forcefield have also given a reasonable description of the MDM2 lid IDR dynamics.^38,39,47^

Estimation of a free-energy surface for the lid region was carried out using a MD simulation-sampling protocol that combines accelerated molecular dynamics (aMD),^48^ Umbrella Sampling (US),^49^ and variational free-energy profile (vFEP) methodologies (see SI text, Table S2, Figure S5 for details).^50^ This approach has been used in previous studies of MDM2 lid dynamics,^38,39^ but ordered lid conformations could not be detected in the computed apo ensembles. Evaluation of the energetics of ordered lid conformation require by use of a free energy surface require finding a low-dimensionality projection that separates ordered and disordered lid conformations. This was achieved here by carrying out independent aMD simulations on models of apo-MDM2, Nutlin-3a/MDM2 and AM-7209/MDM2 initiated with the lid IDR in distinct conformations observed in NMR and X-ray experiments. Snaphots sampled from the different aMD simulations were pooled and projected on different 2D and 3D collective variables drawn from geometric descriptors supported by the AMBER16 suite for follow up US simulations. To avoid excessively time-consuming calculations, we settled down on a 2D representation that monitors the end-to-end distance of the lid and the relative position of the lid with respect to the core region (Figure 3A). Such representation was able to separate ordered lid conformations observed in the AM-7209 complex from other conformations seen during the aMD simulations. Analysis of the free-energy surface obtained after vFEP processing of the US trajectories indicates that in apo MDM2 the lid IDR adopts a major ‘‘closed and disordered’’ lid macrostate that accounts for ca. 75-80% of the lid’s conformational ensemble (Table S3). This macrostate is characterised by a diverse range of lid conformations that extend above helix h2 and the p53-binding site, with occasional formation of helical motifs in the middle of the IDR region (Fig 3B-D). Formation of an ‘‘open and disordered’’ macrostate is energetically strongly disfavoured, and it accounts for less than 1% of the ensemble in the present simulations (Fig 3B-D). An ‘‘open and ordered’’’ macrostate featuring comparatively lower positional fluctuations throughout the lid IDR and a helix-turn-strand motif, similar to that seen in X-ray structures of AM-7209/MDM2 complexes, accounts for ca. 6-8% of the ensemble (Fig 3B-C). Such a low population may explain why it would have been difficult to observe directly the ‘‘open and ordered’’ lid macrostate in previous NMR experiments performed on apo MDM2.^37^

**Figure 3.**
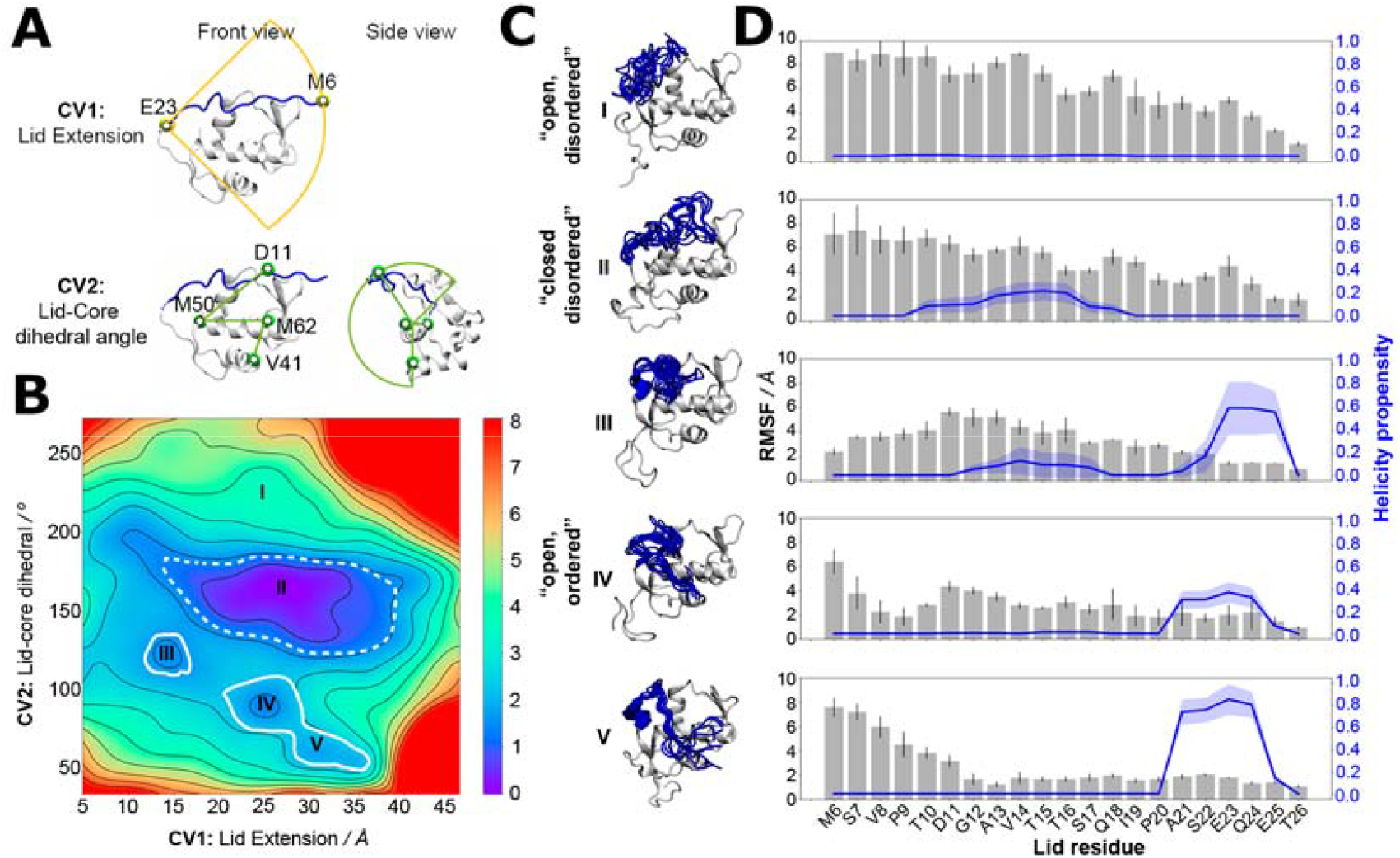
The lid IDR predominantly adopts a closed and disordered macrostate in apo MDM2. **A)** Collective variables used for aMD/US/vFEP calculations. **B)** Free-energy surface of apo MDM2. Heatmap scale in kcal.mol^-1^. Contours used to estimate macrostate populations are shown with solid lines (open and ordered macrostate) or dashed lines (closed and disordered macrostate). **C)** Representative snapshots from different macrostates highlighted in the free-energy surface are shown with the lid IDR highlighted in blue. **D)** Structural properties of the lid in the distinct states highlighted in panels B and C. Error bars denote ±1σ from the first and last halves of the US simulations.

### An adaptive sampling protocol enables efficient computation of binding energetics for MDM2 ligands

The free energy surface analysis indicates that the ‘‘open and ordered’’ lid macrostate is less stable than the ‘‘closed and disordered’’ macrostate by ca. 1.4±0.2 kcal.mol^-1^. To clarify why such a macrostate is predominant in the AM-7209/MDM2 complex it is necessary to determine ligand-binding energetics to distinct lid macrostates. Alchemical absolute binding free-energy calculations (ABFE) may in principle be used for this purpose. Here we use a double decoupling methodology with orientational restraints and standard state corrections to compute standard free energies of binding (Figure S6 and SI text).^51–56^

Achieving reasonable convergence of the computed binding free energies with ABFE methodologies remains very challenging for lead-like molecules bound to flexible protein regions. Here we addressed this issue by developing an adaptive sampling protocol that optimises resource allocation to produce free-energy of binding estimates at a fraction of the computing costs required for a standard protocol (Fig 4A). The adaptive protocol was initially benchmarked on the structurally related but simpler ligand, Pip-2, in complex with MDM2-Lid/short. Reliable binding free-energy estimates could be obtained with a standard protocol where the same sampling time of 50-ns is allocated to each of the windows used to decouple the ligand in the bound and the free state (12 for decoupling electrostatic interactions, 26 for decoupling Lennard-Jones interactions in each bound/free stages, giving a total of 76 windows for the full *λ*-schedule). Five replicates are carried out to estimate a mean binding free energy and confidence interval, which amounts to a cumulative sampling time of 19 μs (Fig 4B). Fig 4C indicates that fluctuations in the mean free energy changes between successive windows are unevenly distributed along the *λ*-schedule. A few windows require significantly more sampling time to decrease statistical fluctuations to a level where the overall binding free energy estimate reaches a plateau. These windows correspond to stages of the *λ*-schedule where the bound state ligand Lennard-Jones interactions are partially decoupled. Such states give noisy free energy changes unless long sampling time are used because diffusion of water molecules in the protein binding site is frustrated by the presence of the partially decoupled ligand.

**Figure 4.**
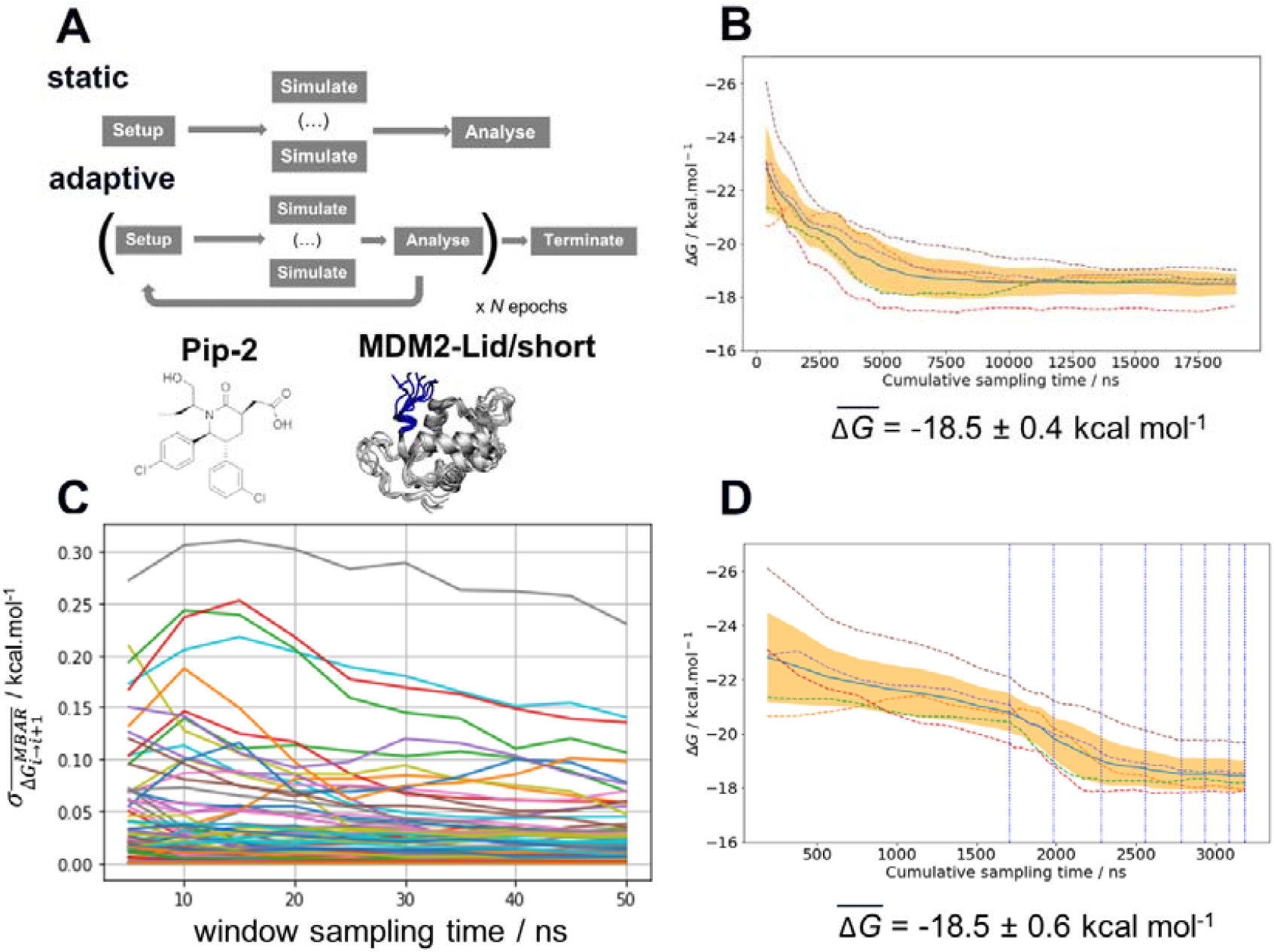
Adaptive sampling of the λ-schedule significantly improves the efficiency of alchemical absolute binding free-energy calculations. **A)** Schematic description of static and adaptive alchemical absolute binding free energy calculation protocols. **B)** Convergence of computed binding free energies for Pip-2 in complex with MDM2-Lid/short with a static protocol. Dashed lines are for individual replicates (*n*=5). The solid blue line denotes the mean of the five replicates and the orange shaded area denotes the 95% confidence interval on the mean of the five replicates. Standard-state corrections have not been applied. **C)** Evolution of the standard deviation of the mean MBAR free-energy change (*n*=5) over time for successive pairs of windows *i, i*+1 across the λ-schedule. Data for each window pair plotted with a different colour for illustrative purposes only. **D)** Convergence profile for free-energies of Pip-2 binding using an adaptive protocol and threshold, τ = 0.100 kcal.mol^-1^. Vertical blue dotted lines denote successive epochs of the adaptive protocol. Other details similar to panel B.

To address this sampling difficulty whilst minimising use of computing resources we implement an adaptive sampling protocol. Quintuplicate simulations are carried out for a period of 5 ns (an ‘‘epoch’’) across a predefined λ-schedule. Once an epoch has completed, the free-energy change between neighbouring windows is estimated using the Multistate Bennett Acceptance Ratio method (MBAR).^57^ Windows whose standard deviation of the mean free-energy change exceeds a threshold parameter, τ, are automatically carried forward to the next epoch. The protocol is iterated for a set number of epochs or until the standard deviation of all windows drops below τ. Thus, as the simulations progress, computing resources are automatically focussed on parts of the *λ*-schedule that require more effort to compute sufficiently precise free energy changes.

Significant savings of compute resource may therefore be achieved by the early termination of sampling of well-behaved windows. Figure 4D depicts a convergence profile for an adaptive sampling run with τ = 0.100 kcal.mol^-1^. The threshold parameter τ must be chosen with care. A too low value yields little savings in computing time, a too high value may cause the simulations to terminate prematurely and yield a binding free energy estimate that deviate from the standard protocol. Additional experiments determined that τ = 0.100 kcal.mol^-1^ gives the best speed/accuracy trade-off for the present system (See SI text and Figure S7). Results statistically indistinguishable from the brute-force calculation are achieved with an almost six-fold decrease in computing resources requirement.

### AM-7209 but not Nutlin-3a shows a marked preference for binding to ordered lid conformations

The efficiency gains observed with the adaptive sampling encouraged us to pursue the more challenging ABFE simulations of AM-7209 and Nutlin-3a in complex with MDM2/Lid-short and MDM2/Lid. For the version with lid, two representative ‘‘closed and disordered’’ and ‘‘open and ordered’’ lid conformations were constructed from the free-energy surface depicted in Figure 3B and previous crystallographic data. The standard free energies of binding computed for the two ligands and the three different MDM2 states are summarised in Figure 5 (Figure S9 for detailed convergence profiles).

**Figure 5.**
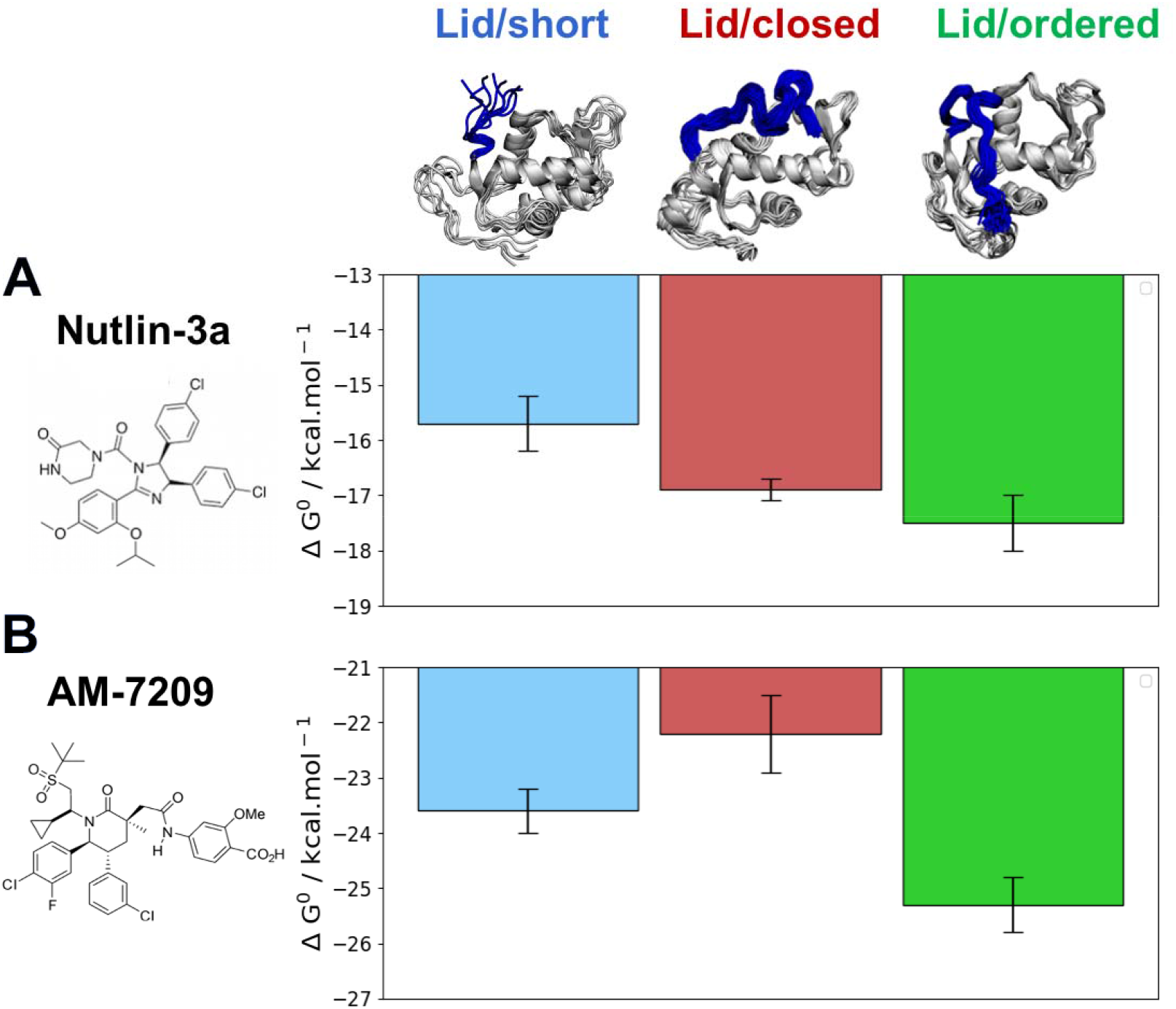
Adaptive ABFE calculations reveal a marked preference of AM-7209 for the MDM2/Lid-closed state. **A)** Computed standard free-energies of binding for Nutlin-3a. **B)** Computed standard binding free-energies for AM-7209. Error bars denote ±1s (*n*=5).

The calculated binding free energies show that Nutlin-3a binds more favourably to MDM2/Lid-closed than MDM2/Lid-short by *ca*. 1 kcal.mol^-1^ (Figure 5A). This is consistent with the slight preference for MDM2/Lid observed in ITC experiments. Nutlin-3a further shows a ca. 0.5 kcal.mol^-1^ preference for binding to MDM2/Lid-ordered when compared to MDM2/Lid-closed. In the case of AM-7209 the MDM2/Lid-closed state is the least preferred, followed by MDM2/Lid-short, and MDM2/Lid-ordered. The latter is favoured over MDM2/Lid-closed by ca. 3 kcal.mol^-1^ and over MDM2/Lid-short by ca. 1.5 kcal.mol^-1^ (Figure 5B).

The overall picture that emerges is that the ‘‘open and ordered’’ lid macrostate of MDM2 is barely detectable in apo MDM2 because it is *ca*. 1.5 kcal.mol^-1^ less stable than the major ‘‘closed and disordered’’ lid microstate. This corresponds to an ‘‘open and ordered’’ equilibrium population of ca. 7%. Upon binding to MDM2 Nutlin-3a shows a slight energetic preference of ca. 0.5 kcal.mol^-1^ for the ‘‘open and ordered’’ lid macrostate, which is insufficient to populate the ordered state to a significant extent (the equilibrium population of ‘‘open and ordered’’ increases to ca. 15%). By contrast the strong (*ca*. 3 kcal.mol^-1^) preference in binding energetics of AM-7209 for the ‘‘open and ordered’’ lid macrostate is sufficient to shift the energetic balance towards significantly populating this macrostate (the equilibrium population of ‘‘open and ordered’’ increases to ca. 90%) in the protein-ligand complex (Figure 6A).

**Figure 6.**
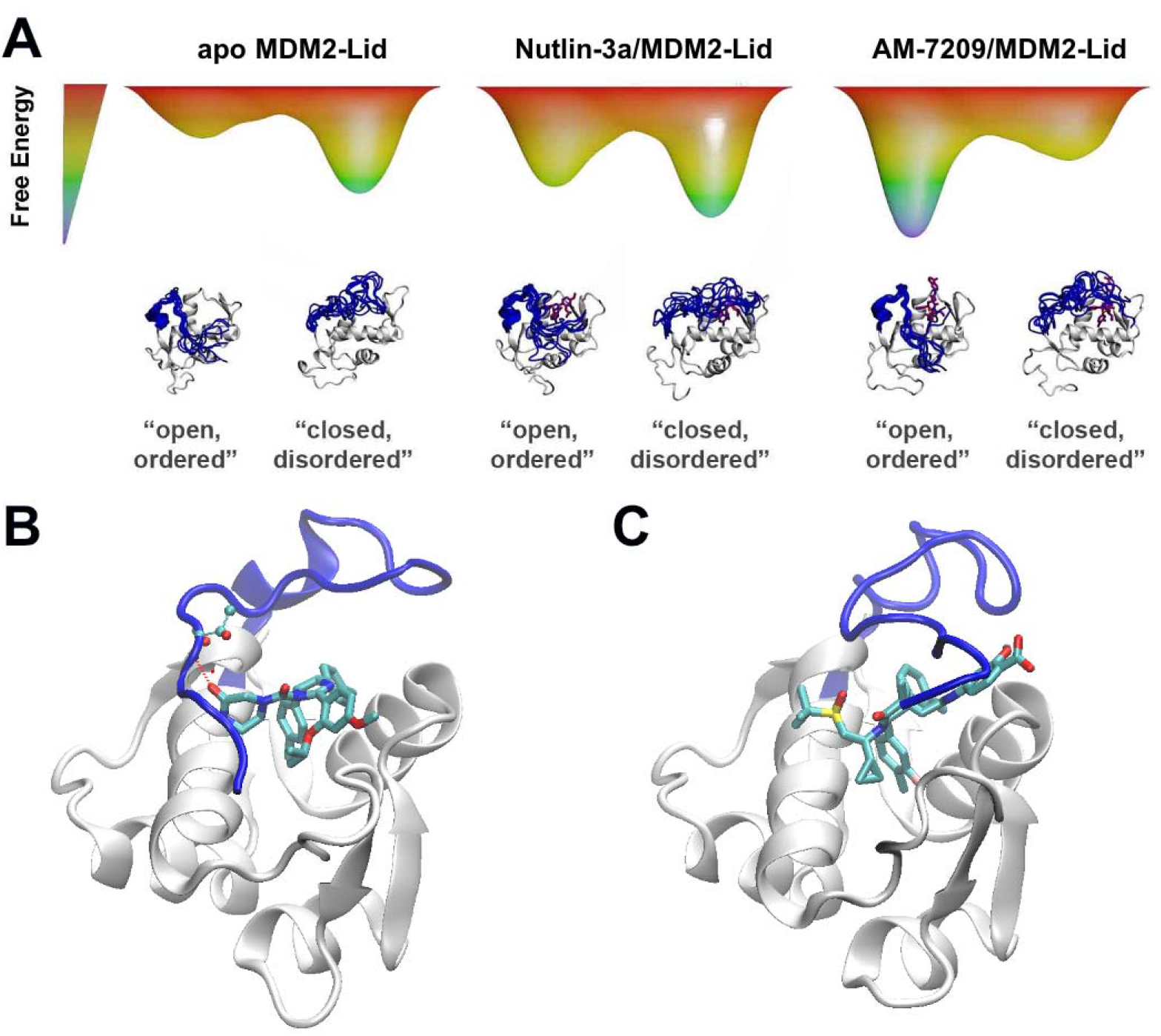
Mechanistic interpretation of ligand-dependent disorder-order transition of the MDM2 lid IDR. **A)** Schematic energy landscapes. **B)** Representative snapshot from simulations of MDM2/Lid-closed in complex with Nutlin-3a. **C)** Representative snapshot from simulations of MDM2/Lid-closed in complex with AM-7209.

In the case of Nutlin-3a interactions with the lid are stabilising as the calculated free-energy of binding to MDM2/Lid-closed is more negative than to MDM2/Lid-short. For AM-7209 the opposite is observed. This difference is not reproduced when using a docking approach, that favours lid-closed states for both ligands (Figure S10). Analysis of the ABFE trajectories suggests that in the ‘‘closed and disordered’’ macrostate the lid lies more frequently above helix *α*2, and the piperazinone group in Nutlin-3a frequently forms hydrogen-bonding interactions with lid residues and in particular the backbone-NH of Thr10 (Figure 6B). By contrast, bulkier ring substituents in AM-7209 hinder interactions of lid residues with helix *α*2 and offer little opportunities for hydrogen-bonding interactions with lid residues (Figure 6C). In apo MDM2, ‘‘closed and disordered’’ lid conformations remain well hydrated by interfacial water molecules or engage in transient hydrogen bonding interactions with MDM2 core residues. This suggests that energetic preferences for the closed lid state may be tuned by varying the nature of contacts between ligands and lid residues. For instance, in the case of Nutlin-3a replacement of the piperazinone solubilising group by a bulkier and less polar moiety projecting over helix *α*2 could be pursued to destabilise interactions of the ‘‘closed and disordered’’ lid state with helix *α*2.

## Conclusions

The present study combined calorimetric analysis of protein-ligand interactions with simulations of molecular dynamics to elucidate the energetics of a small molecule-dependent disorder-to-order protein transition. Such conformational changes are frequently observed in protein-protein interactions yet are rarely exploited in the design of drug-like small molecules. The remarkable 250-fold loss in affinity for AM-7209 arising from truncation of the lid IDR of MDM2 is shown to result from the strong selectivity of this ligand for the ‘‘open and ordered’’ lid macrostate of MDM2 over its ‘‘closed and, disordered’’ microstate. This illustrates the potential value of a ligand-design strategy that seeks to maximise the difference in computed free-energies of binding between ordered and disordered states of an appropriate IDR within the target protein. Such an exercise might be hampered by its demands on computing resources, but the adaptive sampling protocol described here for alchemical absolute calculations of binding free energy affords significant savings in compute time and warrants further methodological investigation. More broadly, the present computational strategy, when applied to other IDRs, could benefit future medicinal chemistry programs that target flexible proteins using small molecules.

## Methods

Proteins were produced by using pET-20b expression vectors hosted by *Escherichia coli*, C41 or C43 (DE3) cells. Nutlin-3a samples were purchased from APExBio. AM-7209 samples were kindly donated by Amgen. ITC measurements were carried out on a Microcal Auto ITC-200 and the data analysed with MicroCal PEAQ-ITC software version 1.1.0.^58^ Accelerated molecular-dynamics and Umbrella sampling simulations were performed using the AMBER16 software suite.^59^ Free-energy surfaces were produced with vFEP 0.1.^60^ Alchemical free energy calculations were prepared using utilities from the software FESetup,^61^ and AMBER16 release, and executed using the SOMD software,^62^ as available in Sire release 2019.1^63^ linked to OpenMM 7.3.1.^64^ Detailed experimental and computational protocol details are provided in the Supporting Information.

## Acknowledgements

J. M. was supported by a Royal Society University Research Fellowship. C. M. M. was supported by a Newton international fellowship. Gratitude is expressed to Amgen for providing materials for this research via its extramural research program. We thank Peter Gimeson from Malvern Panalytical for helpful discussions about ITC measurements. The research leading to these results has received funding from the European Research Council under the European Seventh Framework Programme (FP7/2007-2013)/ERC grant agreement No. 336289. This project made use of time on the ARCHER UK National Supercomputing Service (http://www.archer.ac.uk) granted via the UK High-End Computing Consortium for Biomolecular Simulation, HECBioSim (http://hecbiosim.ac.uk), supported by EPSRC (grant no. EP/L000253/1), and by the Edinburgh Parallel Computing Centre. This work was supported by Wellcome Trust Multi-User Equipment grant 101527/Z13/Z.

## Graphical TOC

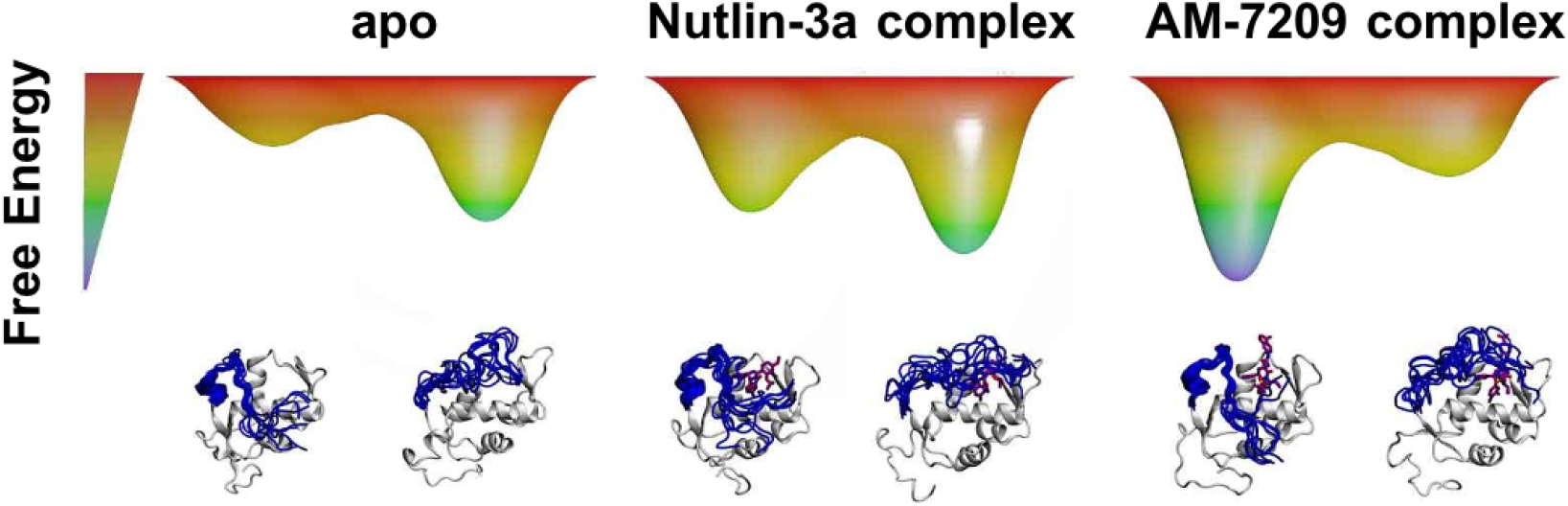

## References

(1) Burger, V. M.; Gurry, T.; Stultz, C. M. Intrinsically Disordered Proteins: Where Computation Meets Experiment. Polymers (Basel). 2014, 6 (10), 2684–2719. https://doi.org/10.3390/polym6102684.

(2) Basile, W.; Salvatore, M.; Bassot, C.; Elofsson, A. Why Do Eukaryotic Proteins Contain More Intrinsically Disordered Regions? PLOS Comput. Biol. 2019, 15 (7), 1–20. https://doi.org/10.1371/journal.pcbi.1007186.

(3) Santofimia-Castaño, P.; Rizzuti, B.; Xia, Y.; Abian, O.; Peng, L.; Velázquez-Campoy, A.; Neira, J. L.; Iovanna, J. Targeting Intrinsically Disordered Proteins Involved in Cancer. Cell. Mol. Life Sci. 2020, 77 (9), 1695–1707. https://doi.org/10.1007/s00018-019-03347-3.

(4) Mollica, L.; Bessa, L. M.; Hanoulle, X.; Jensen, M. R.; Blackledge, M.; Schneider, R. Binding Mechanisms of Intrinsically Disordered Proteins: Theory, Simulation, and Experiment. Frontiers in Molecular Biosciences. 2016, p 52.

(5) Hammoudeh, D. I.; Follis, A. V.; Prochownik, E. V; Metallo, S. J. Multiple Independent Binding Sites for Small-Molecule Inhibitors on the Oncoprotein c-Myc. J. Am. Chem. Soc. 2009, 131 (21), 7390–7401. https://doi.org/10.1021/ja900616b.

(6) Leach, B. I.; Kuntimaddi, A.; Schmidt, C. R.; Cierpicki, T.; Johnson, S. A.; Bushweller, J. H. Leukemia Fusion Target AF9 Is an Intrinsically Disordered Transcriptional Regulator That Recruits Multiple Partners via Coupled Folding and Binding. Structure 2013, 21 (1), 176–183. https://doi.org/10.1016/J.STR.2012.11.011.

(7) Neira, J. L.; Bintz, J.; Arruebo, M.; Rizzuti, B.; Bonacci, T.; Vega, S.; Lanas, A.; Velázquez-Campoy, A.; Iovanna, J. L.; Abián, O. Identification of a Drug Targeting an Intrinsically Disordered Protein Involved in Pancreatic Adenocarcinoma. Sci. Rep. 2017, 7 (1), 39732. https://doi.org/10.1038/srep39732.

(8) Baker, J. M.R.; Hudson, R. P.; Kanelis, V.; Choy, W.-Y.; Thibodeau, P. H.; Thomas, P. J.; Forman-Kay, J. D. CFTR Regulatory Region Interacts with NBD1 Predominantly via Multiple Transient Helices. Nat. Struct. Mol. Biol. 2007, 14 (8), 738–745. https://doi.org/10.1038/nsmb1278.

(9) Milles, S.; Mercadante, D.; Aramburu, I. V.; Jensen, M. R.; Banterle, N.; Koehler, C.; Tyagi, S.; Clarke, J.; Shammas, S. L.; Blackledge, M.; Gräter, F.; Lemke, E. A. Plasticity of an Ultrafast Interaction between Nucleoporins and Nuclear Transport Receptors. Cell 2015, 163 (3), 734–745. https://doi.org/10.1016/j.cell.2015.09.047.

(10) Borgia, A.; Borgia, M. B.; Bugge, K.; Kissling, V. M.; Heidarsson, P. O.; Fernandes, C. B.; Sottini, A.; Soranno, A.; Buholzer, K. J.; Nettels, D.; Kragelund, B. B.; Best, R. B.; Schuler, B. Extreme Disorder in an Ultrahigh-Affinity Protein Complex. Nature 2018, 555 (7694), 61–66. https://doi.org/10.1038/nature25762.

(11) Papadakos, G.; Sharma, A.; Lancaster, L. E.; Bowen, R.; Kaminska, R.; Leech, A. P.; Walker, D.; Redfield, C.; Kleanthous, C. Consequences of Inducing Intrinsic Disorder in a High-Affinity Protein–Protein Interaction. J. Am. Chem. Soc. 2015, 137 (16), 5252–5255. https://doi.org/10.1021/ja512607r.

(12) Rogers, J. M.; Wong, C. T.; Clarke, J. Coupled Folding and Binding of the Disordered Protein PUMA Does Not Require Particular Residual Structure. J. Am. Chem. Soc. 2014, 136 (14), 5197–5200. https://doi.org/10.1021/ja4125065.

(13) Martinelli, A. H. S.; Lopes, F. C.; John, E. B. O.; Carlini, C. R.; Ligabue-Braun, R. Modulation of Disordered Proteins with a Focus on Neurodegenerative Diseases and Other Pathologies. Int. J. Mol. Sci. 2019, 20 (6). https://doi.org/10.3390/ijms20061322.

(14) Lu, H.; Zhou, Q.; He, J.; Jiang, Z.; Peng, C.; Tong, R.; Shi, J. Recent Advances in the Development of Protein–Protein Interactions Modulators: Mechanisms and Clinical Trials. Signal Transduct. Target. Ther. 2020, 5 (1), 213. https://doi.org/10.1038/s41392-020-00315-3.

(15) Heller, G. T.; Aprile, F. A.; Michaels, T. C. T.; Limbocker, R.; Perni, M.; Ruggeri, F. S.; Mannini, B.; Löhr, T.; Bonomi, M.; Camilloni, C.; De Simone, A.; Felli, I. C.; Pierattelli, R.; Knowles, T. P. J.; Dobson, C. M.; Vendruscolo, M. Small-Molecule Sequestration of Amyloid-β as a Drug Discovery Strategy for Alzheimer’s Disease. Sci. Adv. 2020, 6 (45), eabb5924. https://doi.org/10.1126/sciadv.abb5924.

(16) Wallin, S. Intrinsically Disordered Proteins: Structural and Functional Dynamics. Res. Rep. Biol. 2017, Volume 8, 7–16. https://doi.org/10.2147/RRB.S57282.

(17) Cuchillo, R.; Michel, J. Mechanisms of Small-Molecule Binding to Intrinsically Disordered Proteins. Biochem. Soc. Trans. 2012, 40 (5), 1004–1008. https://doi.org/10.1042/BST20120086.

(18) Robustelli, P.; Piana, S.; Shaw, D. E. Mechanism of Coupled Folding-upon-Binding of an Intrinsically Disordered Protein. J. Am. Chem. Soc. 2020, 142 (25), 11092–11101. https://doi.org/10.1021/jacs.0c03217.

(19) Sormanni, P.; Piovesan, D.; Heller, G. T.; Bonomi, M.; Kukic, P.; Camilloni, C.; Fuxreiter, M.; Dosztanyi, Z.; Pappu, R. V; Babu, M. M.; Longhi, S.; Tompa, P.; Dunker, A. K.; Uversky, V. N.; Tosatto, S. C. E.; Vendruscolo, M. Simultaneous Quantification of Protein Order and Disorder. Nat. Chem. Biol. 2017, 13 (4), 339–342. https://doi.org/10.1038/nchembio.2331.

(20) Mu, J.; Liu, H.; Zhang, J.; Luo, R.; Chen, H.-F. Recent Force Field Strategies for Intrinsically Disordered Proteins. J. Chem. Inf. Model. 2021, 61 (3), 1037–1047. https://doi.org/10.1021/acs.jcim.0c01175.

(21) Abian, O.; Vega, S.; Neira, J. L.; Velazquez-Campoy, A. Chapter 17 - High-Throughput Screening for Intrinsically Disordered Proteins by Using Biophysical Methods. In Protein Homeostasis Diseases; Pey, A. L., Ed.; Academic Press, 2020; pp 359–387. https://doi.org/10.1016/B978-0-12-819132-3.00017-8.

(22) Michel, J.; Cuchillo, R. The Impact of Small Molecule Binding on the Energy Landscape of the Intrinsically Disordered Protein C-Myc. PLoS One 2012, 7 (7), e41070.

(23) Heller, G. T.; Sormanni, P.; Vendruscolo, M. Targeting Disordered Proteins with Small Molecules Using Entropy. Trends Biochem. Sci. 2015, 40 (9), 491–496. https://doi.org/10.1016/j.tibs.2015.07.004.

(24) Ruan, H.; Sun, Q.; Zhang, W.; Liu, Y.; Lai, L. Targeting Intrinsically Disordered Proteins at the Edge of Chaos. Drug Discov. Today 2019, 24 (1), 217–227. https://doi.org/10.1016/j.drudis.2018.09.017.

(25) Robustelli, P.; Piana, S.; Shaw, D. E. Developing a Molecular Dynamics Force Field for Both Folded and Disordered Protein States. Proc. Natl. Acad. Sci. 2018, 115 (21), E4758 LP–E4766. https://doi.org/10.1073/pnas.1800690115.

(26) Robustelli, P.; Ibanez-de-Opakua, A.; Campbell-Bezat, C.; Giordanetto, F.; Becker, S.; Zweckstetter, M.; Pan, A. C.; Shaw, D. E. Molecular Basis of Small-Molecule Binding to α-Synuclein. bioRxiv 2021, 2021.01.22.426549. https://doi.org/10.1101/2021.01.22.426549.

(27) Nag, S.; Qin, J.; Srivenugopal, K. S.; Wang, M.; Zhang, R. The MDM2-P53 Pathway Revisited. J. Biomed. Res. 2013, 27 (4), 254–271. https://doi.org/10.7555/JBR.27.20130030.

(28) Demir, Ö.; Barros, E. P.; Offutt, T. L.; Rosenfeld, M.; Amaro, R. E. An Integrated View of P53 Dynamics, Function, and Reactivation. Curr. Opin. Struct. Biol. 2021, 67, 187–194. https://doi.org/10.1016/j.sbi.2020.11.005.

(29) Ding, K.; Lu, Y.; Nikolovska-Coleska, Z.; Qiu, S.; Ding, Y.; Gao, W.; Stuckey, J.; Krajewski, K.; Roller, P. P.; Tomita, Y.; Parrish, D. A.; Deschamps, J. R.; Wang, S. Structure-Based Design of Potent Non-Peptide MDM2 Inhibitors. J. Am. Chem. Soc. 2005, 127 (29), 10130–10131. https://doi.org/10.1021/ja051147z.

(30) Zhao, Y.; Liu, L.; Sun, W.; Lu, J.; McEachern, D.; Li, X.; Yu, S.; Bernard, D.; Ochsenbein, P.; Ferey, V.; Carry, J.-C.; Deschamps, J. R.; Sun, D.; Wang, S. Diastereomeric Spirooxindoles as Highly Potent and Efficacious MDM2 Inhibitors. J. Am. Chem. Soc. 2013, 135 (19), 7223–7234. https://doi.org/10.1021/ja3125417.

(31) Hansen, M. J.; Feringa, F. M.; Kobauri, P.; Szymanski, W.; Medema, R. H.; Feringa, B. L. Photoactivation of MDM2 Inhibitors: Controlling Protein–Protein Interaction with Light. J. Am. Chem. Soc. 2018, 140 (41), 13136–13141. https://doi.org/10.1021/jacs.8b04870.

(32) Konopleva, M.; Martinelli, G.; Daver, N.; Papayannidis, C.; Wei, A.; Higgins, B.; Ott, M.; Mascarenhas, J.; Andreeff, M. MDM2 Inhibition: An Important Step Forward in Cancer Therapy. Leukemia 2020, 34 (11), 2858–2874. https://doi.org/10.1038/s41375-020-0949-z.

(33) Wang, W.; Qin, J.-J.; Rajaei, M.; Li, X.; Yu, X.; Hunt, C.; Zhang, R. Targeting MDM2 for Novel Molecular Therapy: Beyond Oncology. Med. Res. Rev. 2020, 40 (3), 856–880. https://doi.org/10.1002/med.21637.

(34) Rew, Y.; Sun, D.; Yan, X.; Beck, H. P.; Canon, J.; Chen, A.; Duquette, J.; Eksterowicz, J.; Fox, B. M.; Fu, J.; Gonzalez, A. Z.; Houze, J.; Huang, X.; Jiang, M.; Jin, L.; Li, Y.; Li, Z.; Ling, Y.; Lo, M.-C.; Long, A. M.; McGee, L. R.; McIntosh, J.; Oliner, J. D.; Osgood, T.; Saiki, A. Y.; Shaffer, P.; Wang, Y. C.; Wortman, S.; Yakowec, P.; Ye, Q.; Yu, D.; Zhao, X.; Zhou, J.; Medina, J. C.; Olson, S. H. Discovery of AM-7209, a Potent and Selective 4-Amidobenzoic Acid Inhibitor of the MDM2–P53 Interaction. J. Med. Chem. 2014, 57 (24), 10499–10511. https://doi.org/10.1021/jm501550p.

(35) Michelsen, K.; Jordan, J. B.; Lewis, J.; Long, A. M.; Yang, E.; Rew, Y.; Zhou, J.; Yakowec, P.; Schnier, P. D.; Huang, X.; Poppe, L. Ordering of the N-Terminus of Human MDM2 by Small Molecule Inhibitors. J. Am. Chem. Soc. 2012, 134 (41), 17059–17067. https://doi.org/10.1021/ja305839b.

(36) Vassilev, L. T.; Vu, B. T.; Graves, B.; Carvajal, D.; Podlaski, F.; Filipovic, Z.; Kong, N.; Kammlott, U.; Lukacs, C.; Klein, C.; Fotouhi, N.; Liu, E. A. In Vivo Activation of the P53 Pathway by Small-Molecule Antagonists of MDM2. Science (80-.). 2004, 303 (5659), 844–848. https://doi.org/10.1126/science.1092472.

(37) Showalter, S. A.; Bruschweiler-Li, L.; Johnson, E.; Zhang, F.; Brüschweiler, R. Quantitative Lid Dynamics of MDM2 Reveals Differential Ligand Binding Modes of the P53-Binding Cleft. J. Am. Chem. Soc. 2008, 130 (20), 6472–6478. https://doi.org/10.1021/ja800201j.

(38) Bueren-Calabuig, J. A.; Michel, J. Impact of Ser17 Phosphorylation on the Conformational Dynamics of the Oncoprotein MDM2. Biochemistry 2016, 55 (17), 2500–2509. https://doi.org/10.1021/acs.biochem.6b00127.

(39) Bueren-Calabuig, J. A.; Michel, J. Elucidation of Ligand-Dependent Modulation of Disorder-Order Transitions in the Oncoprotein MDM2. PLOS Comput. Biol. 2015, 11 (6), 1–27. https://doi.org/10.1371/journal.pcbi.1004282.

(40) Song, D.; Luo, R.; Chen, H.-F. The IDP-Specific Force Field Ff14IDPSFF Improves the Conformer Sampling of Intrinsically Disordered Proteins. J. Chem. Inf. Model. 2017, 57 (5), 1166–1178. https://doi.org/10.1021/acs.jcim.7b00135.

(41) Shabane, P. S.; Izadi, S.; Onufriev, A. V. General Purpose Water Model Can Improve Atomistic Simulations of Intrinsically Disordered Proteins. J. Chem. Theory Comput. 2019, 15 (4), 2620–2634. https://doi.org/10.1021/acs.jctc.8b01123.

(42) Somavarapu, A. K.; Kepp, K. P. The Dependence of Amyloid-β Dynamics on Protein Force Fields and Water Models. ChemPhysChem 2015, 16 (15), 3278– 3289. https://doi.org/10.1002/cphc.201500415.

(43) Man, V. H.; He, X.; Derreumaux, P.; Ji, B.; Xie, X.-Q.; Nguyen, P. H.; Wang, J. Effects of All-Atom Molecular Mechanics Force Fields on Amyloid Peptide Assembly: The Case of Aβ16–22 Dimer. J. Chem. Theory Comput. 2019, 15 (2), 1440–1452. https://doi.org/10.1021/acs.jctc.8b01107.

(44) Zapletal, V.; Mládek, A.; Melková, K.; Louša, P.; Nomilner, E.; Jaseňáková, Z.; Kubáň, V.; Makovická, M.; Laníková, A.; Žídek, L.; Hritz, J. Choice of Force Field for Proteins Containing Structured and Intrinsically Disordered Regions. Biophys. J. 2020, 118 (7), 1621–1633. https://doi.org/10.1016/j.bpj.2020.02.019.

(45) Gao, Y.; Zhang, C.; Wang, X.; Zhu, T. A Test of AMBER Force Fields in Predicting the Secondary Structure of α-Helical and β-Hairpin Peptides. Chem. Phys. Lett. 2017, 679, 112–118. https://doi.org/10.1016/j.cplett.2017.04.074.

(46) Zheng, Y.; Cui, Q. The Histone H3 N-Terminal Tail: A Computational Analysis of the Free Energy Landscape and Kinetics. Phys. Chem. Chem. Phys. 2015, 17 (20), 13689–13698. https://doi.org/10.1039/C5CP01858G.

(47) Mukherjee, S.; Pantelopulos, G. A.; Voelz, V. A. Markov Models of the Apo-MDM2 Lid Region Reveal Diffuse yet Two-State Binding Dynamics and Receptor Poses for Computational Docking. Sci. Rep. 2016, 6 (1), 31631. https://doi.org/10.1038/srep31631.

(48) Pierce, L. C. T.; Salomon-Ferrer, R.; Augusto F. De Oliveira, C.; McCammon, J. A.; Walker, R. C. Routine Access to Millisecond Time Scale Events with Accelerated Molecular Dynamics. J. Chem. Theory Comput. 2012, 8 (9), 2997– 3002. https://doi.org/10.1021/ct300284c.

(49) Torrie, G. M.; Valleau, J. P. Nonphysical Sampling Distributions in Monte Carlo Free-Energy Estimation: Umbrella Sampling. J. Comput. Phys. 1977, 23 (2), 187–199. https://doi.org/10.1016/0021-9991(77)90121-8.

(50) Lee, T.-S.; Radak, B. K.; Huang, M.; Wong, K.-Y.; York, D. M. Roadmaps through Free Energy Landscapes Calculated Using the Multidimensional VFEP Approach. J. Chem. Theory Comput. 2014, 10 (1), 24–34. https://doi.org/10.1021/ct400691f.

(51) Jorgensen, W. L.; Buckner, J. K.; Boudon, S.; Tirado[Rives, J. Efficient Computation of Absolute Free Energies of Binding by Computer Simulations. Application to the Methane Dimer in Water. J. Chem. Phys. 1988, 89 (6), 3742–3746. https://doi.org/10.1063/1.454895.

(52) Gilson, M. K.; Given, J. A.; Bush, B. L.; McCammon, J. A. The Statistical-Thermodynamic Basis for Computation of Binding Affinities: A Critical Review. Biophys. J. 1997, 72 (3), 1047–1069. https://doi.org/10.1016/S0006-3495(97)78756-3.

(53) Aldeghi, M.; Heifetz, A.; Bodkin, M. J.; Knapp, S.; Biggin, P. C. Predictions of Ligand Selectivity from Absolute Binding Free Energy Calculations. J. Am. Chem. Soc. 2017, 139 (2), 946–957. https://doi.org/10.1021/jacs.6b11467.

(54) Aldeghi, M.; Heifetz, A.; Bodkin, M. J.; Knapp, S.; Biggin, P. C. Accurate Calculation of the Absolute Free Energy of Binding for Drug Molecules. Chem. Sci. 2016, 7 (1), 207–218. https://doi.org/10.1039/C5SC02678D.

(55) Mey, A. S. J. S.; Allen, B. K.; Macdonald, H. E. B.; Chodera, J. D.; Hahn, D. F.; Kuhn, M.; Michel, J.; Mobley, D. L.; Naden, L. N.; Prasad, S.; Rizzi, A.; Scheen, J.; Shirts, M. R.; Tresadern, G.; Xu, H. Best Practices for Alchemical Free Energy Calculations [Article v1.0]. Living J. Comput. Mol. Sci. 2020, 2 (1), 18378. https://doi.org/10.33011/livecoms.2.1.18378.

(56) Cournia, Z.; Allen, B. K.; Beuming, T.; Pearlman, D. A.; Radak, B. K.; Sherman, W. Rigorous Free Energy Simulations in Virtual Screening. J. Chem. Inf. Model. 2020, 60 (9), 4153–4169. https://doi.org/10.1021/acs.jcim.0c00116.

(57) Shirts, M. R.; Chodera, J. D. Statistically Optimal Analysis of Samples from Multiple Equilibrium States. J. Chem. Phys. 2008, 129 (12), 124105. https://doi.org/10.1063/1.2978177.

(58) Malvern inc. MicroCal PEAQ-ITC 1.1.0.

(59) Case, D. A.; Betz, R. M.; Cerutti, D. S.; Cheatham III, T. E.; Darden, T. A.; Duke, R. E.; Giese, T. J.; Gohlke, H.; Goetz, A. W.; Homeyer, N.; Izadi, S.; Janowski, P.; Kaus, J.; Kovalenko, A.; Lee, T. S.; LeGrand, S.; Li, P.; Lin, C.; Luchko, T.; Luo, R.; Madej, B.; Mermelstein, D.; Merz, K. M.; Monard, G.; Nguyen, H.; Nguyen, H. T.; Omelyan, I.; Onufriev, A.; Roe, D. R.; Roitberg, A.; Sagui, C.; Simmerling, C. L.; Botello-Smith, W. M.; Swails, J.; Walker, R. C.; Wang, J.; Wolf, R. M.; Wu, X.; Xiao, L.; Kollman, P. A. Amber 16. University of California, San Francisco. 2016. https://doi.org/10.1002/jcc.23031.

(60) Lee, T.-S.; Radak, B. K.; Pabis, A.; York, D. M. A New Maximum Likelihood Approach for Free Energy Profile Construction from Molecular Simulations. J. Chem. Theory Comput. 2013, 9 (1), 153–164. https://doi.org/10.1021/ct300703z.

(61) Loeffler, H. H.; Michel, J.; Woods, C. FESetup: Automating Setup for Alchemical Free Energy Simulations. J. Chem. Inf. Model. 2015, 55 (12), 2485– 2490. https://doi.org/10.1021/acs.jcim.5b00368.

(62) Calabrò, G.; Woods, C. J.; Powlesland, F.; Mey, A. S. J. S.; Mulholland, A. J.; Michel, J. Elucidation of Nonadditive Effects in Protein–Ligand Binding Energies: Thrombin as a Case Study. J. Phys. Chem. B 2016, 120 (24), 5340– 5350. https://doi.org/10.1021/acs.jpcb.6b03296.

(63) Woods, C.; Mey, A. S. J. S.; Calabrò, G.; Michel, J. Sire Molecular Simulation Framework. 2019.

(64) Eastman, P.; Swails, J.; Chodera, J. D.; McGibbon, R. T.; Zhao, Y.; Beauchamp, K. A.; Wang, L.-P.; Simmonett, A. C.; Harrigan, M. P.; Stern, C. D.; Wiewiora, R. P.; Brooks, B. R.; Pande, V. S. OpenMM 7: Rapid Development of High Performance Algorithms for Molecular Dynamics. PLOS Comput. Biol. 2017, 13 (7), e1005659.

