## Supporting Information for "Energetics of a protein disorder-order transition in small molecule recognition"

**Table of Contents**


| | |
|---|---|
| **Methods text** | S3 |
| **SI references** | S13 |
| **Figures and Tables** | S15 |

#### Methods

##### Protein samples preparation.

Plasmid constructs were ordered from commercial GeneArt gene synthesis services. The specific gene sequence of human MDM2-Lid (6-125 aa.) and human MDM2-Lid/short (17-125 aa.) were inserted in a pET-20b expression vector (ampicillin resistant), with a C-terminal hexahistidine-tag included after the protein sequence. The primary sequence for MDM2-Lid was: MSVPTDGAVTTSQIPASEQETLVRPKPLLLKLLKSVGAQKDTYTMKEVLFYLGQYIMTK RLYDEKQQHIVYCSNDLLGDLFGVPSFSVKEHRKIYTMIYRNLVVVNQQESSDSGTSVS ENHHHHHH. The primary sequence for MDM2-Lid/short was: SQIPASEQETLVRPKPLLLKLLKSVGAQKDTYTMKEVLFYLGQYIMTKRLYDEKQQHIV YCSNDLLGDLFGVPSFSVKEHRKIYTMIYRNLVVVNQQESSDSGTSVSENHHHHHH. Proteins were overproduced in *Escherichia coli,* C41 (DE3) for the MDM2-Lid and C43 (DE3) host cells for MDM2-Lid/short, respectively. Bacterial cultures grown overnight in LB broth medium supplemented with carbenicillin (100 mg/mL) were induced with 1.0 M isopropyl-β-D-1-thiogalactopyranoside (IPTG) after reaching $O.D._{600}$ ~1.0 and then further incubated at 25 °C for 4-6 hours. Cells were harvested and lysed in lysis buffer (20 mM Tris, 100 mM NaCl, 1 mM tris-(2-carboxyethyl) phosphine -TCEP-, pH 8.0) using a Constant Cell Systems Disruptor (1.1 kW TS Benchtop) set at 25 kpsi and a repeat cycle of lysis was further performed. The lysed fraction was centrifuged, and the resultant supernatant directly applied to a $Ni^{2+}$-affinity (GE Healthcare) column. Elution of C-terminal tagged protein was achieved with a stepwise (8% then 70%) gradient of 20 mM Tris, 100 mM NaCl, 100 mM imidazole, pH 8.0 buffer. The final protein purification was achieved by passing concentrated samples through a 16/60 Superdex75 size-exclusion chromatography (GE Healthcare) pre-equilibrated with 50 mM phosphate, 100 mM NaCl, 1 mM TCEP, pH 7.0. Identify of the proteins was confirmed by LC-MS (**Figure S1** and **Figure S2**), which showed that the MDM2 residues sequence in the purified proteins was 7-125 for MDM2/Lid and 17-125 for MDM2-Lid/short.

**Calorimetric measurements**

ITC was used to measure the binding affinity ($K_d$) of the MDM2 ligands Nutlin-3a and AM-7209. Nutlin-3a was purchased from commercial vendor APExBIO, whereas AM-7209 samples were kindly provided by Amgen inc. The buffer used for all ITC experiments was 50 mM phosphate, 100 mM NaCl, 1 mM TCEP, pH 7.0. All titrations were performed using a MicroCal Auto-iTC200 isothermal titration calorimeter from Malvern. Analytical and non-linear curve fitting of integrated raw data were carried out assuming one site of binding in the MicroCal PEAQ-ITC Analysis Software version 1.1.0. Titration experiments for every protein construct were made by using a first injection of 0.4 µl followed by 19 injections of 2 µl, with an interval between each injection of 300 sec. The temperature in the cell was 25 ºC and sample stirring in the cell was kept at 750 rpm. The concentration of the protein in the cell was 10 µM, and for ligands in the syringe 100 µM Nutlin-3a or 150 µM AM-7209. Throughout all titrations, the final DMSO concentration was maintained at 1 % (v/v) for the cell and the syringe. For the titration of MDM2-Lid with AM-7209, the binding affinity was too strong to be measured by a direct titration, hence a competitive-titration protocol was employed. For this protocol, a solution of MDM2-Lid (10 µM) and Nutlin-3a (100 µM) was incubated for two hours and then the pre-formed MDM2-Lid/-Nutlin-3a complex was titrated with 150 µM of AM-7209 in the syringe. The binding affinity was calculated using the protocol of Sigurskjold et al, as implemented in MicroCal PEAQ-ITC.[1] Representative titrations for the two ligands are shown in **Figure S3** and **Figure S4**. Values for the thermodynamic measurements reported in Figure 2 of the manuscript are shown in **Table S1**. Our calculated $K_d$ values for Nutlin-3a to MDM2-Lid and MDM2-Lid/short are *ca.* three-fold smaller than that reported by Michelsen et al. This difference could arise from the higher (5% v/v) amount of DMSO used in titrations by Michelsen *et al.*, as DMSO has been shown in several cases to compete with ligand for access to protein surfaces.[2] In addition, the MDM2/6-125 protein construct used by Michelsen also contained four extra amino acids after cleavage of the N-terminal fusion tag, whereas our constructs contained a hexahistidine tag at the C terminus of MDM2 to minimise impact on the conformational preference of the lid IDR.

**Accelerated Molecular Dynamics (aMD) simulations**

*Models preparation*

Initial exploration of the conformational landscape of MDM2 lid was performed by means of accelerated molecular dynamics (aMD) simulations.[3] As starting points, we used three different MDM2 (residues 6-125) structures that differ in the conformation of the lid: semi-open state (obtained from model 2 of the NMR ensemble PDB id 1Z1M[4]), closed state and ordered state reported by Bueren-Calabuig et al.[5] Missing residues of the C-terminal domain were modelled based on model 2 of the NMR ensemble PDB id 2LZG (residues 1-125).[6] The protein and ligand complexes were solvated in a triclinic TIP3P water box with edges extending 18 Å away from the edges of the protein,[7] and $Cl^-$ anions were added to neutralize the system. The ff99SBildnnmr force field was used to model the protein,[8] and Joung and Cheatham parameters were used for the anions.[9] Nutlin-3a and AM-7209 ligands were parameterized using GAFF atom types,[10,11] and atomic charges derived with the AM1-BCC method,[12] as implemented in the *antechamber* tool of the AMBER16 suite.[13]

Each system was energy minimized with 4500 steps of steepest descent followed by 4500 steps of conjugate gradient and subsequently smoothly heated from 100 K to 298 K over a 1 ns-long simulation at constant volume, with positional restraints on protein and ligand atoms. Then each system was subjected to a 4-ns equilibration simulation, with restraints being gradually reduced for protein and ligand. Finally, the density of the system was equilibrated by means of a 1-ns long simulation at 298 K in the NPT ensemble. Throughout the heating and equilibration stages, SHAKE was used for all bonds involving hydrogen atoms,[14] and a time step of 2 fs was employed. Prior to the aMD simulations, a conventional 100-ns long simulation in the NPT ensemble was performed using the GPU accelerated version of PMEMD from the AMBER16 software package.

*Accelerated Molecular Dynamics simulation parameters*

aMD adds a positive energy boost to the potential energy function allowing the simulated systems to efficiently explore different regions of the potential energy surface (PES) through reduced energy barriers. A dual boost approach was used, with a potential energy boost applied to all the heavy atoms and an additional energy boost applied to all the torsion angles in the system.

To prevent an intense energy boost from pushing the system towards high energy regions of the PES that may lead to unrealistic behaviour, such as protein unfolding, residues in the MDM2 core region (25-125) were restrained using harmonic restraints with a force constant of 20 kcal mol$^{-1}$ A$^{-2}$. For each system, 300-ns long aMD trajectories were obtained.

Initial $E_P$, $E_D$, $\alpha_P$ and $\alpha_D$ parameters for the aMD runs were selected following the guidelines of Pierce and co-workers .[3] Trial runs were carried out to tune the parameters to achieve broad sampling of lid conformations. The values of $E_P$, $E_D$, $\alpha_P$ and $\alpha_D$ described in **Table S2** were adopted.

*Collective variables definition*

Two suitable collective variables (CV) were selected to describe the conformational preferences of the MDM2 lid as observed in the aMD simulations. CV1 defines the extension of the lid as the distance between the Cα atoms of residues M6 and E23. CV2 is the dihedral angle defined by the Cα atoms of residues D11, M50, M62 and V41 (lid-core dihedral angle). Figure 3A in the manuscript depicts both CV.

The conformational space (CV space) defined by these two collective variables allowed us to discriminate four different states for the lid conformation: open (CV2 > 220º), semi-open (CV1 < 24 Å, 110º < CV2 < 220º), closed (CV1 > 24 Å, 110º < CV2 < 220º) and ordered conformations (CV2 < 110º).

*Umbrella sampling protocol*

An US protocol was devised to compute the equilibrium distribution of the lid conformational ensembles. The CV space was sampled within the CV1 values 5 - 45 Å with an interval of 2 Å and CV2 values 36 - 268 º with an interval of 8 º accounting for a total of 524 bins. The initial coordinates were selected by choosing the closest snapshot to the average structure sampled for that bin during the preceding aMD runs. The selected protein structures were extracted and used as the initial starting conformations. The protein was solvated again following the protocol described previously. Prior to the US runs, the starting conformations were re-equilibrated using the same energy minimisation protocol and a brief equilibration protocol (500 ps of thermalisation and 500 ps equilibration in the NPT ensemble). Then, 4-ns long production runs were performed using the same conditions described in the system setup section. Monitoring of the convergence profile of the FES along each CV suggested this was sufficient to obtain reasonably converged FES (**Figure S5**). Harmonic potential restraints of 1 kcal $mol^{-1}Å^{-2}$ and 0.12 kcal $mol^{-1}deg^{-2}$ were applied to the target CV1 and CV2 values respectively. These small restraining force constants were chosen to avoid excessive energetic penalties for conformations slightly deviating from the target CV values. Values of the reaction coordinates were stored every 10 fs for post-processing.

*2D variational free-energy profile (vFEP) analysis*

The 2D variational vFEP method was used to obtain unbiased free energy profiles along the defined CV space.[15] vFEP is a maximum likelihood parametric approach to reweight biased simulation data. For smoothly varying free-energy surfaces, the method has been shown to yield converged FES with fewer windows and with a fraction of the statistics required with the Weighted Histogram Analysis Method. To estimate uncertainties in free energies, all US trajectories were sub-divided into two parts of equal duration (2 ns) and analysed separately (**Table S3**).

Inputs files, scripts and instructions to repeat the aMD/US/vFEP calculations are available at https://github.com/michellab/MDM2-DG_paper.

**Alchemical absolute binding free energy calculations**

*Preparation of MDM2/ligands input files for free energy calculations*

The protein-ligand structures for the free-energy calculations were obtained from the X-ray crystal structures with PDB IDs: 4HG7 for Nutlin-3a/MDM2,[16] and 4WT2 for AM-7209/MDM2.[17] All water molecules were removed from the structures and all proteins were capped at the C terminus and N terminus with N- methyl and acetyl groups respectively. The coordinates of the structured lid conformation for the three protein-ligand complexes were taken from the Pip-2/MDM2 crystal structure,[6] while the coordinates of the "closed" lid conformation were extracted from the calculated FES.

Input files for the free-energy simulations were created using utilities from the AMBER16 software suite,[13] and the software FESetup.[18] Protein parameterisation was performed using the ff14SB Amber force-field,[19] while ligands were parameterised using the GAFF2 forcefield,[10,11] and AM1-BCC partial charges.[12] The system was solvated in a cubic box with TIP3P water molecules,[7] with a minimum distance between the protein and the edges of the box of 12 Å. Counter ions were added to neutralize the total net charge. The same approach was followed for parameterising the ligand in the free phase.

Next an equilibration protocol was applied to relax the box size. Initially, energy minimization of the entire system was implemented with 1000 steps of steepest descent gradients, using the software sander. Then, an NVT protocol was followed for 200 ps at 298 K, followed by an NPT equilibration for a further 200 ps at 1 atm. Finally, a 2-ns NPT MD simulation was run with the software SOMD to reach a final density of about 1 g $cm^{-3}$.[20,21] The final coordinate files were retrieved using the software cpptraj.

*Adaptive sampling protocol*

Alchemical free energy simulations were performed following a double decoupling protocol.[22,23] In this method, a ligand $L$ is mutated into a "non-interacting" molecule both in the solvated and the bound phase using a two-step process. An example of the thermodynamic cycle used in these calculations is illustrated in **Figure S6**.

During the protein-bound simulations the ligand is restrained to prevent drifting out of the binding site and to accelerate convergence of the free energy changes. A standard state correction term $\Delta G^{o}_{rest}$ is applied post-simulation to work out the free-energy change upon release of the ligand restraints to standard state condition. Details of the restraint protocol used for these simulations are given below.

The overall cycle gives a standard free-energy of binding as shown by eq 1:

$$(1)\ \Delta G^{o} = (\Delta G_{host}^{q=0} + \Delta G_{host}^{vdW=0}) - (\Delta G_{solv}^{q=0} + \Delta G_{solv}^{vdW=0}) + \Delta G_{rest}^{o}$$

For the MDM2-ligand complexes, both bound and free phase *discharging* steps were run with twelve $\lambda$ windows (0.000, 0.050, 0.100, 0.200, 0.300, 0.400, 0.500, 0.600, 0.700, 0.800, 0.900, 1.000), while 26 λ windows (0.000, 0.025, 0.050, 0.075, 0.100, 0.125, 0.150, 0.200, 0.250, 0.300, 0.350, 0.400, 0.450, 0.500, 0.550, 0.600, 0.650, 0.700, 0.750, 0.800, 0.850, 0.900, 0.950, 0.970, 0.990, 1.000) were employed for the *vanishing* step, both in complex and solvated phase.

The adaptive sampling approach involves running an initial set of calculations in which each $\lambda$ window is simulated for a duration of 5 ns with SOMD in the NPT ensemble. Temperature control is maintained with an Andersen Thermostat with a coupling constant of 10 $ps^{-1}$.[24] Pressure control is achieved by a Monte Carlo barostat that attempts isotropic box-edge scaling every 100 fs. A 12 Å atom-based cutoff distance for the non-bonded interactions is used, using a Barker Watts reaction field with dielectric constant of 78.3.[25]

Free-energy changes were estimated with the multistate Bennet acceptance ratio method as implemented in the Sire utility *analyse_freenrg*.[26] To achieve a more robust estimation of free energies, each simulation was repeated five times, using different initial velocities drawn from the Maxwell-Boltzmann distribution and statistical uncertainties are reported as 95% of the standard error of the mean.

For the adaptive sampling protocol, the simulations were carried out initially for a duration of 5 ns. Following this, we identify $\lambda$ windows that contribute the most to the overall uncertainty of the absolute binding free-energy by calculating the standard deviation of the free-energy estimates between neighbouring MBAR windows. Only simulations whose uncertainty exceeds a threshold

$\tau$ are carried further into the next iteration of the protocol. **Figure S7** shows that the optimal choice of t is around 0.1 kcal.mol$^{-1}$ for the benchmarked system because lower values yield similar results with additional computing costs. Higher values yield more time savings, but the free energy estimate deviates from the reference results.

*Restraints protocol*

The choice of the restraints is important for these calculations as it influences the convergence of the final binding free energy. For instance, the ligands used to inhibit MDM2, due to their large size, can adopt multiple orientations during the simulations once their interactions with the surrounding environment are weakened. This translates into an increase in computing time requirements to sample all the thermally accessible orientations and thus slows down convergence of the final free energy of binding. To reduce the sampling time needed for these calculations, we need to prevent the ligand from tumbling and drifting away from the host cavity. For this purpose, a series of flat-bottom distance restraints were defined between four ligand atoms and a variable number of protein atoms, depending on the MDM2/ligand complexes. Details of restraint parameters are given in **Figure S8**. Unless otherwise noted the width of the flat-bottom region was $D_{ji}$ = 2 Å and a half-harmonic penalty of $k_{ji}$= 10 kcal mol$^{-1}$Å$^{-2}$ was applied for distances deviating from the flat bottom. The free-energy change for releasing the restraints and bring the decoupled ligands to standard state concentration and free rotation was determined by numerical integration as described in Bosisio *et al*.[27] and the resulting values are given in **Table S3**.

The convergence profile of the free energies of binding of AM-7209 and Nutlin-3a bound to different lid states is shown in **Figure S9**. Additional epochs were run until the mean binding free energy estimates were deemed stable. In comparison with Pip2/MDM2-short the results are noisier for simulations including the long lid versions, but the binding selectivity trend in binding energies of AM-7209 between MDM2/Lid-closed and MDM2/Lid-ordered state appears robust.

Inputs files, scripts and instructions to repeat the ABFE calculations are available at https://github.com/michellab/MDM2-DG_paper.

*Control docking study.*

The equilibrated conformations of the protein ligand complexes used at the start of the ABFE calculations were loaded in the software Flare 5.[28] The ''scoring in place with optimisation'' mode from the Docking module of Flare was used to estimate a DG score. The results are shown in **Figure S11**. Similar trends were observed with other metrics such as VS Score. The results suggest that the scoring function used in Flare 5 overestimates binding energies to the MDM2/Lid-closed state. This may be because this state forms more direct contact with the ligands.

### References


(1) Sigurskjold, B. W. Exact Analysis of Competition Ligand Binding by Displacement Isothermal Titration Calorimetry. *Anal. Biochem.* **2000**, *277* (2), 260–266. https://doi.org/10.1006/abio.1999.4402.

(2) Cubrilovic, D.; Zenobi, R. Influence of Dimehylsulfoxide on Protein–Ligand Binding Affinities. *Anal. Chem.* **2013**, *85* (5), 2724–2730. https://doi.org/10.1021/ac303197p.

(3) Pierce, L. C. T.; Salomon-Ferrer, R.; Augusto F. De Oliveira, C.; McCammon, J. A.; Walker, R. C. Routine Access to Millisecond Time Scale Events with Accelerated Molecular Dynamics. *J. Chem. Theory Comput.* **2012**, *8* (9), 2997–3002. https://doi.org/10.1021/ct300284c.

(4) Uhrinova, S.; Uhrin, D.; Powers, H.; Watt, K.; Zheleva, D.; Fischer, P.; McInnes, C.; Barlow, P. N. Structure of Free MDM2 N-Terminal Domain Reveals Conformational Adjustments That Accompany P53-Binding. *J. Mol. Biol.* **2005**, *350* (3), 587–598. https://doi.org/https://doi.org/10.1016/j.jmb.2005.05.010.

(5) Bueren-Calabuig, J. A.; Michel, J. Elucidation of Ligand-Dependent Modulation of Disorder-Order Transitions in the Oncoprotein MDM2. *PLOS Comput. Biol.* **2015**, *11* (6), 1–27. https://doi.org/10.1371/journal.pcbi.1004282.

(6) Michelsen, K.; Jordan, J. B.; Lewis, J.; Long, A. M.; Yang, E.; Rew, Y.; Zhou, J.; Yakowec, P.; Schnier, P. D.; Huang, X.; Poppe, L. Ordering of the N-Terminus of Human MDM2 by Small Molecule Inhibitors. *J. Am. Chem. Soc.* **2012**, *134* (41), 17059–17067. https://doi.org/10.1021/ja305839b.

(7) Jorgensen, W. L.; Chandrasekhar, J.; Madura, J. D.; Impey, R. W.; Klein, M. L. Comparison of Simple Potential Functions for Simulating Liquid Water. *J. Chem. Phys.* **1983**, *79* (2), 926–935. https://doi.org/10.1063/1.445869.

(8) Li, D.-W.; Brüschweiler, R. NMR-Based Protein Potentials. *Angew. Chemie Int. Ed.* **2010**, *49* (38), 6778–6780. https://doi.org/https://doi.org/10.1002/anie.201001898.

(9) Joung, I. S.; Cheatham, T. E. Determination of Alkali and Halide Monovalent Ion Parameters for Use in Explicitly Solvated Biomolecular Simulations. *J. Phys. Chem. B* **2008**, *112* (30), 9020–9041. https://doi.org/10.1021/jp8001614.

(10) Wang, J.; Wolf, R. M.; Caldwell, J. W.; Kollman, P. A.; Case, D. A. Development and Testing of a General Amber Force Field. *J. Comput. Chem.* **2004**, *25* (9), 1157–1174. https://doi.org/https://doi.org/10.1002/jcc.20035.

(11) Wang, J.; Wang, W.; Kollman, P. A.; Case, D. A. Automatic Atom Type and Bond Type Perception in Molecular Mechanical Calculations. *J. Mol. Graph. Model.* **2006**, *25* (2), 247–260. https://doi.org/https://doi.org/10.1016/j.jmgm.2005.12.005.

(12) Jakalian, A.; Bush, B. L.; Jack, D. B.; Bayly, C. I. Fast, Efficient Generation of High-Quality Atomic Charges. AM1-BCC Model: I. Method. *J. Comput. Chem.* **2000**, *21* (2), 132–146. https://doi.org/https://doi.org/10.1002/(SICI)1096-987X(20000130)21:2<132::AID-JCC5>3.0.CO;2-P.

(13) Case, D. A.; Betz, R. M.; Cerutti, D. S.; Cheatham III, T. E.; Darden, T. A.; Duke, R. E.; Giese, T. J.; Gohlke, H.; Goetz, A. W.; Homeyer, N.; Izadi, S.; Janowski, P.; Kaus, J.; Kovalenko, A.; Lee, T. S.; LeGrand, S.; Li, P.; Lin, C.; Luchko, T.; Luo, R.; Madej, B.; Mermelstein, D.; Merz, K. M.; Monard, G.; Nguyen, H.; Nguyen, H. T.; Omelyan, I.; Onufriev, A.; Roe, D. R.; Roitberg, A.; Sagui, C.; Simmerling, C. L.; Botello-Smith, W. M.; Swails, J.; Walker, R. C.; Wang, J.; Wolf, R. M.; Wu, X.; Xiao, L.; Kollman, P. A. Amber 16. *University of California, San Francisco*. 2016. https://doi.org/10.1002/jcc.23031.

(14) Ryckaert, J.-P.; Ciccotti, G.; Berendsen, H. J. C. Numerical Integration of the Cartesian Equations of Motion of a System with Constraints: Molecular Dynamics of n-Alkanes. *J.*

*Comput. Phys.* **1977**, *23* (3), 327–341. https://doi.org/https://doi.org/10.1016/0021-9991(77)90098-5.
(15) Lee, T.-S.; Radak, B. K.; Pabis, A.; York, D. M. A New Maximum Likelihood Approach for Free Energy Profile Construction from Molecular Simulations. *J. Chem. Theory Comput.* **2013**, *9* (1), 153–164. https://doi.org/10.1021/ct300703z.
(16) Anil, B.; Riedinger, C.; Endicott, J. A.; Noble, M. E. M. The Structure of an MDM2{--}Nutlin-3a Complex Solved by the Use of a Validated MDM2 Surface-Entropy Reduction Mutant. *Acta Crystallogr. Sect. D* **2013**, *69* (8), 1358–1366. https://doi.org/10.1107/S0907444913004459.
(17) Rew, Y.; Sun, D.; Yan, X.; Beck, H. P.; Canon, J.; Chen, A.; Duquette, J.; Eksterowicz, J.; Fox, B. M.; Fu, J.; Gonzalez, A. Z.; Houze, J.; Huang, X.; Jiang, M.; Jin, L.; Li, Y.; Li, Z.; Ling, Y.; Lo, M.-C.; Long, A. M.; McGee, L. R.; McIntosh, J.; Oliner, J. D.; Osgood, T.; Saiki, A. Y.; Shaffer, P.; Wang, Y. C.; Wortman, S.; Yakowec, P.; Ye, Q.; Yu, D.; Zhao, X.; Zhou, J.; Medina, J. C.; Olson, S. H. Discovery of AM-7209, a Potent and Selective 4-Amidobenzoic Acid Inhibitor of the MDM2–P53 Interaction. *J. Med. Chem.* **2014**, *57* (24), 10499–10511. https://doi.org/10.1021/jm501550p.
(18) Loeffler, H. H.; Michel, J.; Woods, C. FESetup: Automating Setup for Alchemical Free Energy Simulations. *J. Chem. Inf. Model.* **2015**, *55* (12), 2485–2490. https://doi.org/10.1021/acs.jcim.5b00368.
(19) Maier, J. A.; Martinez, C.; Kasavajhala, K.; Wickstrom, L.; Hauser, K. E.; Simmerling, C. Ff14SB: Improving the Accuracy of Protein Side Chain and Backbone Parameters from Ff99SB. *J. Chem. Theory Comput.* **2015**, *11* (8), 3696–3713. https://doi.org/10.1021/acs.jctc.5b00255.
(20) Woods, C.; Michel, J. Sire Molecular Simulation Framework, Revision 1786. 2019.
(21) Eastman, P.; Swails, J.; Chodera, J. D.; McGibbon, R. T.; Zhao, Y.; Beauchamp, K. A.; Wang, L.-P.; Simmonett, A. C.; Harrigan, M. P.; Stern, C. D.; Wiewiora, R. P.; Brooks, B. R.; Pande, V. S. OpenMM 7: Rapid Development of High Performance Algorithms for Molecular Dynamics. *PLOS Comput. Biol.* **2017**, *13* (7), e1005659.
(22) Jorgensen, W. L.; Buckner, J. K.; Boudon, S.; Tirado-Rives, J. Efficient Computation of Absolute Free Energies of Binding by Computer Simulations. Application to the Methane Dimer in Water. *J. Chem. Phys.* **1988**, *89* (6), 3742–3746. https://doi.org/10.1063/1.454895.
(23) Gilson, M. K.; Given, J. A.; Bush, B. L.; McCammon, J. A. The Statistical-Thermodynamic Basis for Computation of Binding Affinities: A Critical Review. *Biophys. J.* **1997**, *72* (3), 1047–1069. https://doi.org/https://doi.org/10.1016/S0006-3495(97)78756-3.
(24) Andersen, H. C. Molecular Dynamics Simulations at Constant Pressure and/or Temperature. *J. Chem. Phys.* **1980**, *72* (4), 2384–2393. https://doi.org/10.1063/1.439486.
(25) Tironi, I. G.; Sperb, R.; Smith, P. E.; van Gunsteren, W. F. A Generalized Reaction Field Method for Molecular Dynamics Simulations. *J. Chem. Phys.* **1995**, *102* (13), 5451–5459. https://doi.org/10.1063/1.469273.
(26) Shirts, M. R.; Chodera, J. D. Statistically Optimal Analysis of Samples from Multiple Equilibrium States. *J. Chem. Phys.* **2008**, *129* (12), 124105. https://doi.org/10.1063/1.2978177.
(27) Bosisio, S.; Mey, A. S. J. S.; Michel, J. Blinded Predictions of Host-Guest Standard Free Energies of Binding in the SAMPL5 Challenge. *J. Comput. Aided. Mol. Des.* **2017**, *31* (1), 61–70. https://doi.org/10.1007/s10822-016-9933-0.
(28) Cresset. Flare 5. 2021.

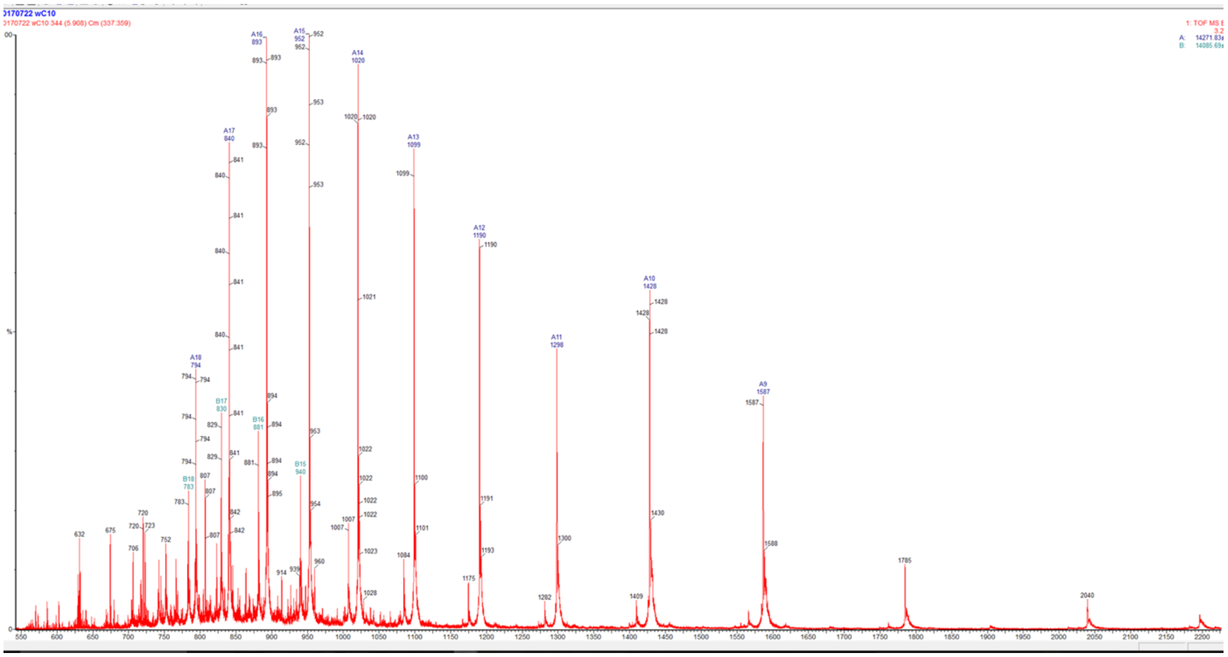

**Figure S1.** Positive mode mass spectrum acquired by LC-MS of MDM2 7-125-His6. The N-terminal methionine was cleaved during purification. The deconvoluted average mass was calculated to be 14271.83 Da, in agreement with the calculated theoretical average mass of 14272.22 Da.

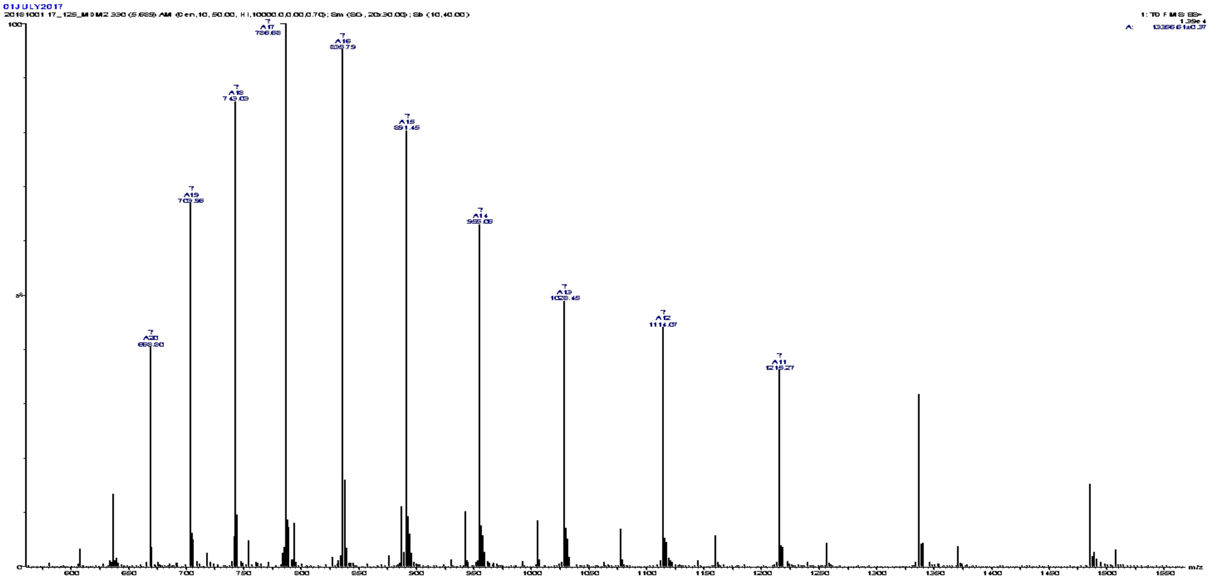

**Figure S2.** Positive mode mass spectrum acquired by LC-MS of MDM2 17-125-His6. Calculated molecular mass for MDM2 17-125 are: [M+H]+ = 13343.22 Da, and [M+Na]+ = 13366.22 Da. The deconvoluted average mass was calculated to be 13366.61 Da, which is consistent with the calculated [M+Na]+ mass.

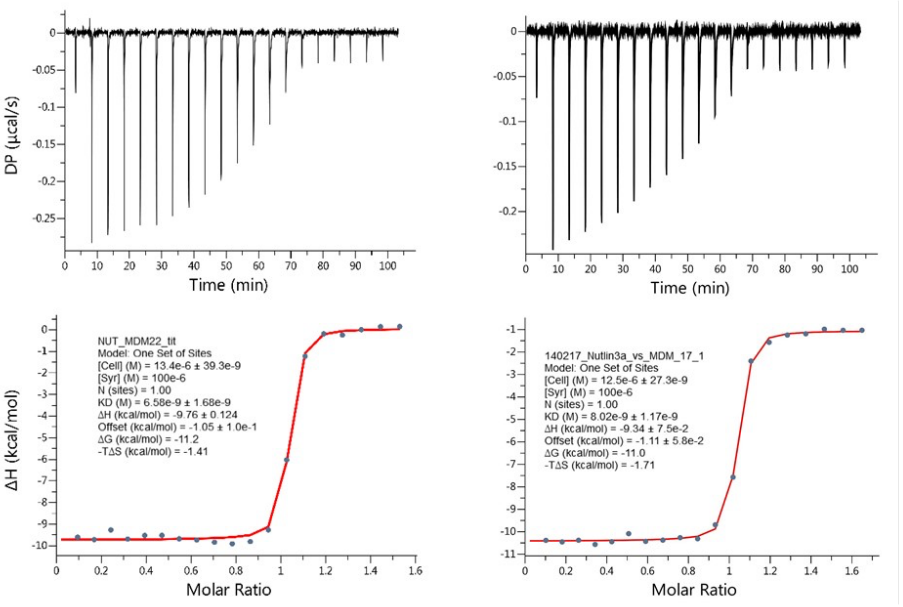


Nutlin-3a/MDM2-Lid Nutlin-3a/MDM2-Lid/short

**Figure S3.** Representative titrations of Nutlin-3a to MDM2-Lid and MDM2-Lid/short constructs.

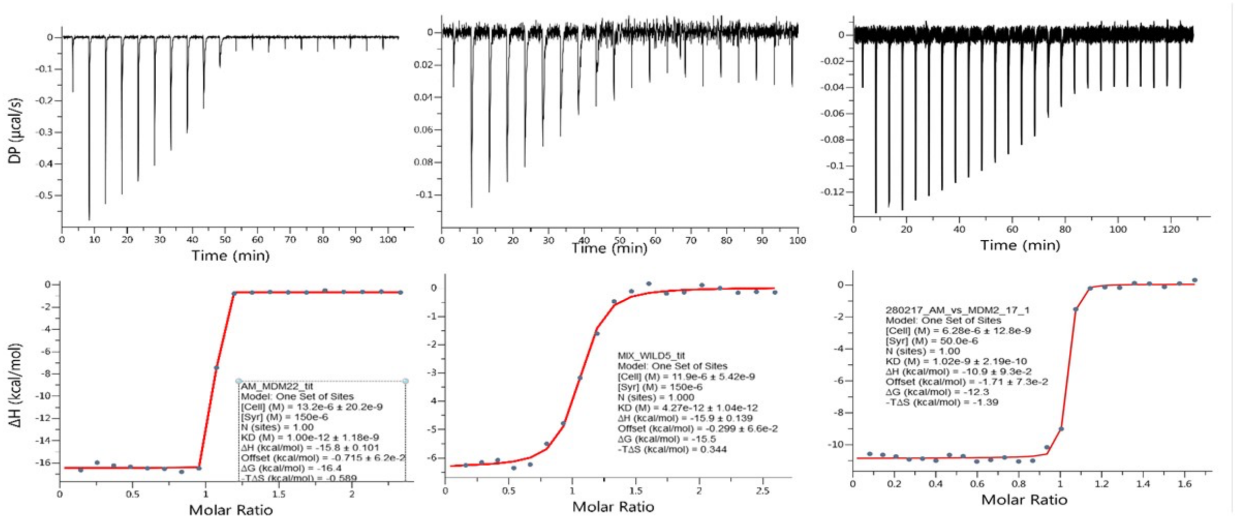


AM-7209/MDM2-Lid Direct titration AM-7209/MDM2-Lid Competitive titration AM-7209/MDM2-Lid/short

**Figure S4.** Representative titrations of AM-7209 to MDM2-Lid (direct and competitive titration) and MDM2-Lid/short constructs.

**Table S1.** Binding thermodynamic parameters from triplicate measurements. Uncertainties are ±1σ.

**Nutlin-3a**

| Construct | $K_d$ / pM | $\Delta G^0$ / kcal.mol$^{-1}$ | $\Delta H$ / kcal.mol$^{-1}$ | $-T\Delta S$ / kcal.mol$^{-1}$ |
|---|---|---|---|---|
| **MDM2-Lid** | 5090 ± 1408 | -11.33 ± 0.16 | -9.48 ± 0.27 | -1.85 ± 0.38 |
| **MDM2-Lid/short** | 6950 ± 1325 | -11.14 ± 0.12 | -9.40 ± 0.42 | -1.73 ± 0.50 |

**AM-7209**

| Construct | $K_d$ / pM | $\Delta G^0$ / kcal.mol$^{-1}$ | $\Delta H$ / kcal.mol$^{-1}$ | $-T\Delta S$ / kcal.mol$^{-1}$ |
|---|---|---|---|---|
| **MDM2-Lid** | 4.1 ± 0.6 | -15.54 ± 0.09 | -16.00 ± 0.22 | 0.44 ± 0.38 |
| **MDM2-Lid/short** | 1040 ± 40 | -12.25 ± 0.03 | -10.90 ± 0.30 | -1.35 ± 0.30 |

**Table S2. Accelerated Molecular Dynamics parameters used in the present work.** The first two terms control the boost applied to the total potential energy, the last two terms control the boost applied to the dihedral energy. Figures in $kcal.mol^{-1}$.

| System | $E_P$ | $\alpha_P$ | $E_D$ | $\alpha_D$ |
|---|---|---|---|---|
| wt-apo | -130310 | 10000 | 3000 | 78 |
| wt-AM7209 | -123360 | 10000 | 3000 | 78 |
| wt-Nutlin-3a | -125038 | 10000 | 3000 | 78 |

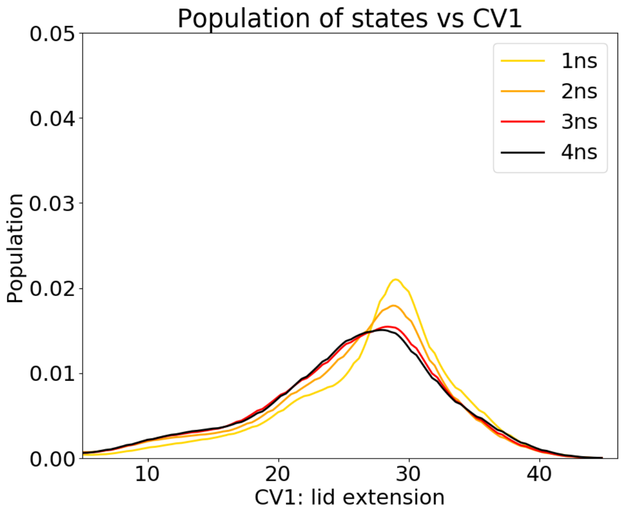


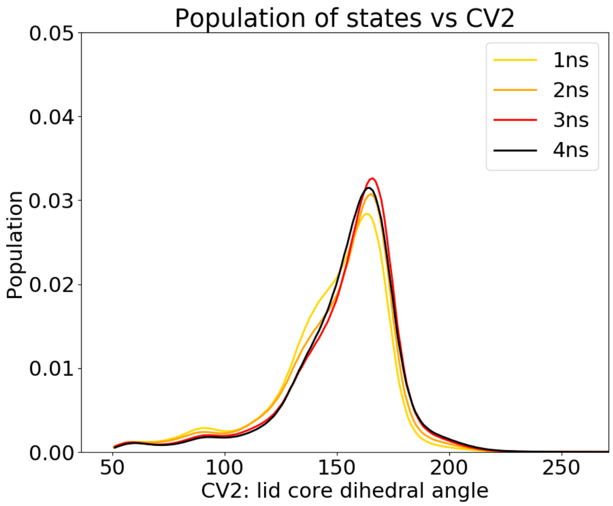


**Figure S5.** Convergence plots for apo MDM2 FES. Populations along each CV value for cumulative sampling time per window of 0-1 ns, 0-2 ns, 0-3 ns, 0-4 ns respectively.

**Table S3.** Population estimates for the areas flagged in Fig 3B.

| | "Closed, disordered" | "Open, ordered" | | |
|---|---|---|---|---|
| **Population (%)** | II | III | IV | V |
| **Full sampling (1-4ns)** | 76.9 | 0.8 | 3.5 | 2.9 |
| **1-2 ns** | 71.1 | 1.2 | 2.7 | 3.4 |
| **3-4 ns** | 70.9 | 0.7 | 5.9 | 3.8 |
| **Mean** | 73.0 | 0.9 | 4.0 | 3.4 |
| **$\sigma$** | 3.5 | 0.3 | 1.6 | 0.5 |

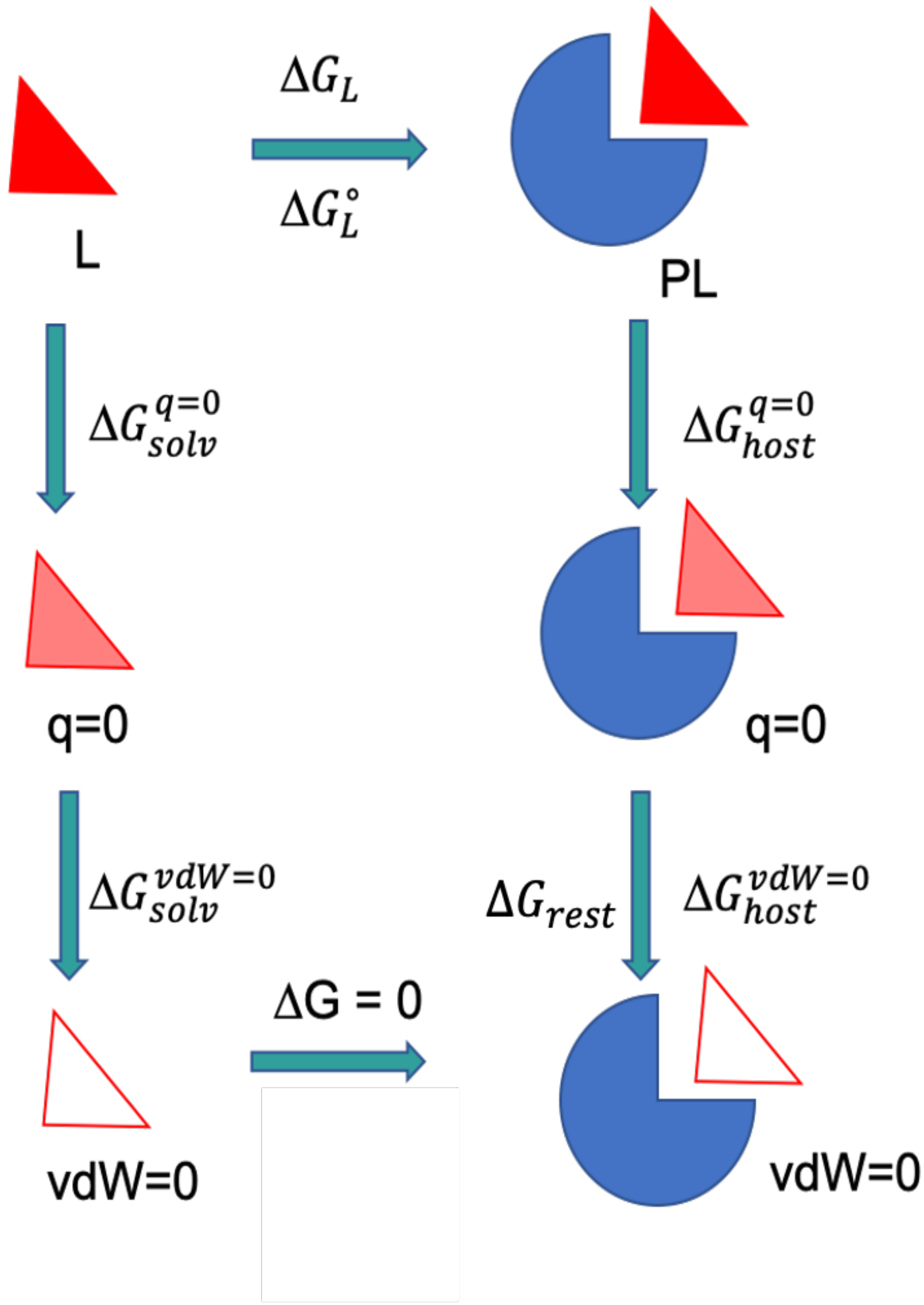


**Figure S6.** An example of the thermodynamic cycle used in absolute free-energy calculations with the double decoupling method. Ligand L is transformed into an ideal thermodynamic state in both solvated and complex phase using a two-step process. In the *discharging* step, partial charges of the ligand are switched off, retrieving free energy changes $\Delta G_{solv}^{q=0}$ and $\Delta G_{host}^{q=0}$. Subsequently, a *vanishing* step is carried on by turning off the vdW terms of the discharged ligand, providing free energy changes $\Delta G_{solv}^{vdW=0}$ and $\Delta G_{host}^{vdW=0}$.

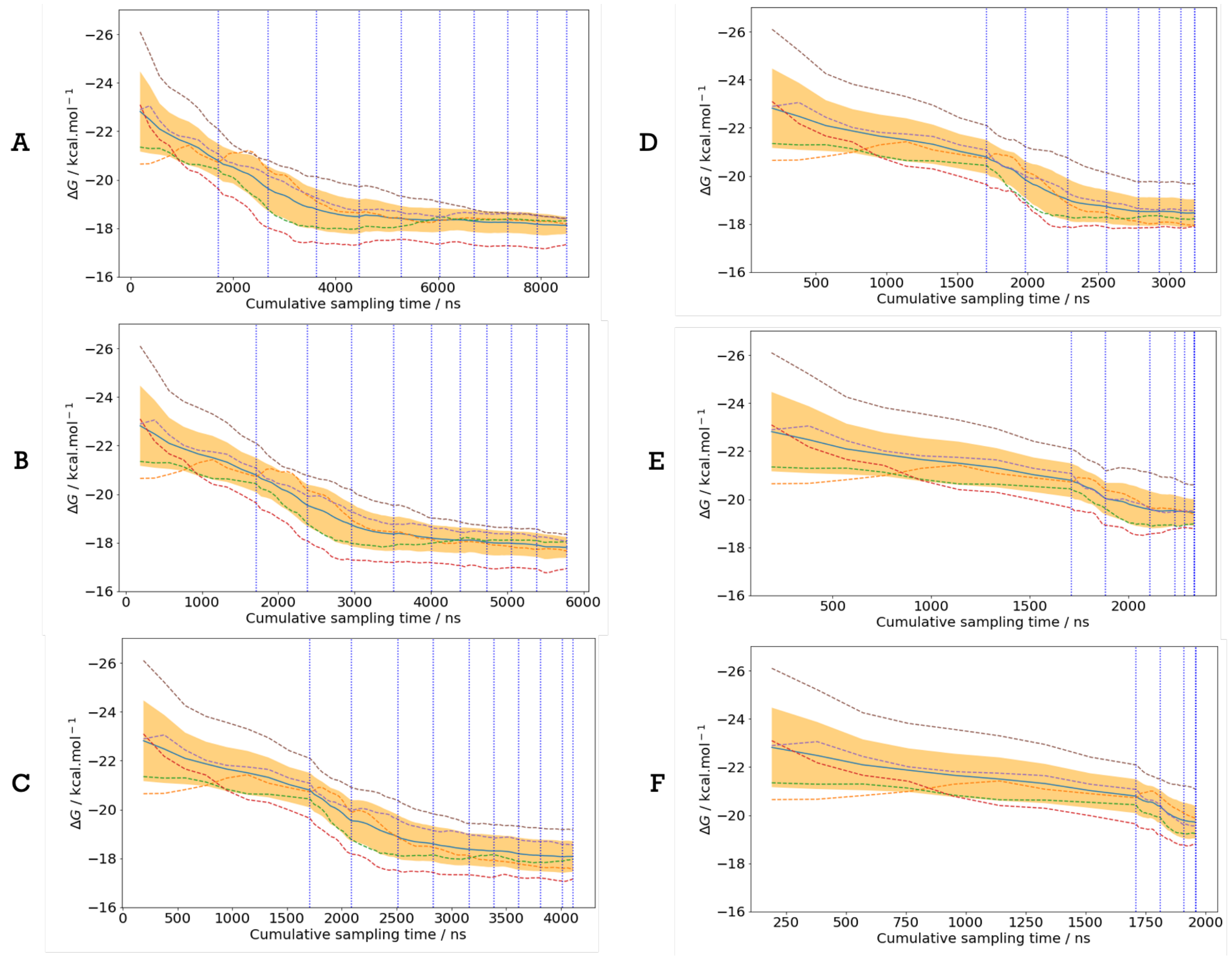


**Figure S7.** Convergence plots of binding free energy of Pip2 bound to MDM2-Lid/short for variable thresholds **A)** $\tau$=0.025 kcal.mol$^{-1}$, **B)** $\tau$=0.050 kcal.mol$^{-1}$ **C)** $\tau$=0.075 kcal.mol$^{-1}$, **D)** $\tau$=0.100 kcal.mol$^{-1}$, **E)** $\tau$=0.150 kcal.mol$^{-1}$, **F)** $\tau$=0.200 kcal.mol$^{-1}$. Formatting scheme identical to Figure 4D of the main manuscript.

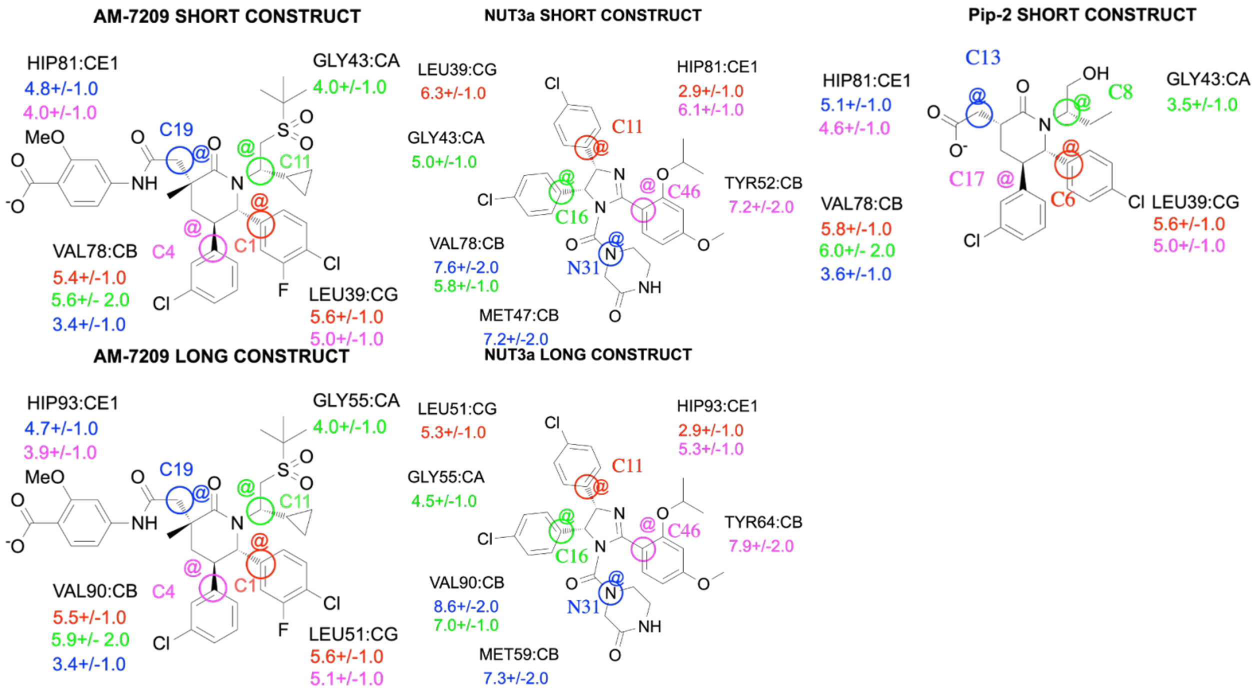


**Figure S8.** Diagram illustrating restraints definition for Pip-2, AM-7209, Nutlin-3a.

**Table S4.** Standard state correction terms and standard binding free energies for the different ABFE simulations reported in this study. All energies in kcal.mol$^{-1}$. Uncertainty given as ±1σ (*n*=5).

| **Ligands** | $\boldsymbol{\Delta G_{restraint}}$ | $\boldsymbol{\Delta G^{o}}$ |
|---|---|---|
| MDM2-Lid/short | | |
| Nutlin-3a | -5.4 ± 0.4 | -15.7 ± 0.5 |
| Pip2 | -5.5 ± 0.2 | -12.9 ± 0.6 |
| AM-7209 | -5.2 ± 0.3 | -23.6 ± 0.4 |
| MDM2-Lid/closed | | |
| Nutlin-3a | -5.3 ± 0.2 | -16.9 ± 0.2 |
| AM-7209 | -5.0 ± 0.2 | -22.2 ± 0.7 |
| MDM2-Lid/ordered | | |
| Nutlin-3a | -5.3 ± 0.2 | -17.5 ± 0.5 |
| AM-7209 | -5.2 ± 0.2 | -25.3 ± 0.5 |

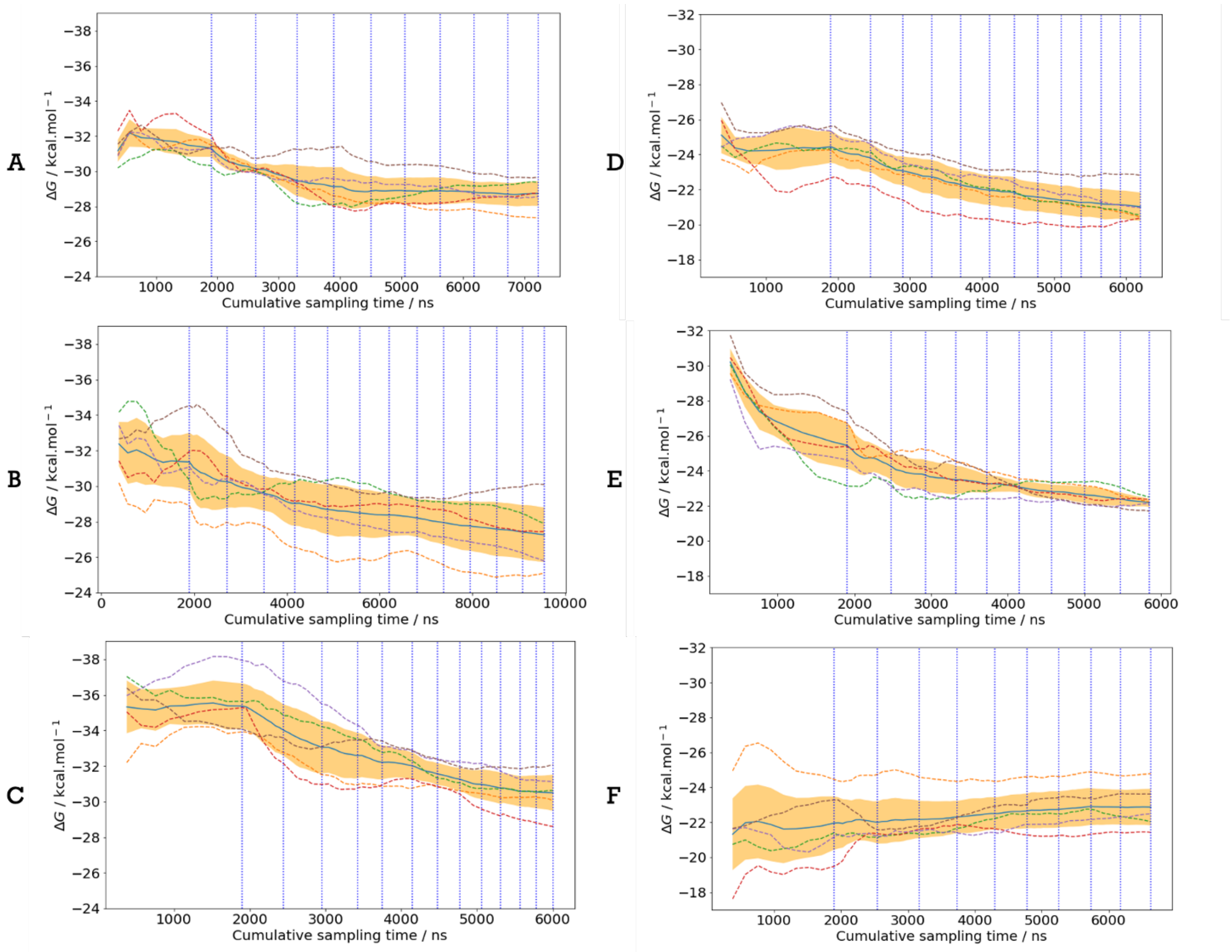


**Figure S9.** 6 Convergence of the free energy of binding ΔG for **A)** AM-7209/MDM2-Lid/short, **B)** AM-7209/MDM2-Lid/closed, **C)** AM-7209/MDM2/Lid-ordered, **D)** Nutlin-3a/MDM2-Lid/short, **E)** Nutlin-3a/MDM2-Lid/closed, **F)** Nutlin-3a/MDM2-Lid/ordered. Formatting scheme identical to Figure 4D of the main manuscript.

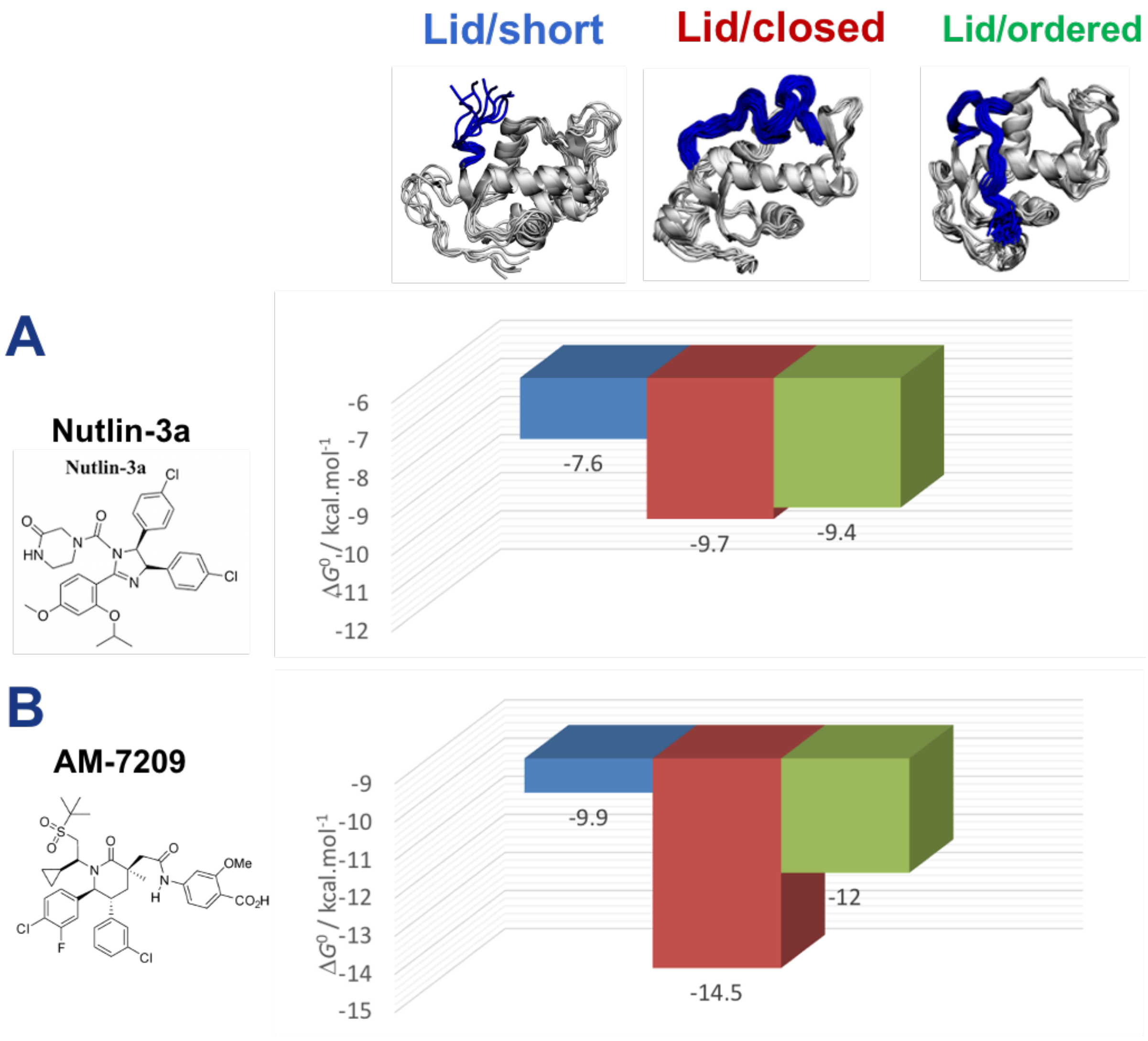


**Figure S10.** Docking energies obtained with Flare 5 for **A)** Nutlin-3a, **B)** AM-7209. Values in kcal.mol$^{-1}$

**Dataset S1.** Input files and scripts to repeat the aMD/US/vFEP and ABFE simulations reported in the manuscript are available at https://github.com/michellab/MDM2-DG_paper

**Movie S1.** Collective variables used for the aMD/US/vFEP calculations. Movie available at https://github.com/michellab/MDM2-DG_paper/blob/main/CVmovie.low.mp4
